# Optogenetic activation of the inhibitory nigro-collicular circuit evokes orienting movements in mice

**DOI:** 10.1101/2020.05.21.107680

**Authors:** C.A. Villalobos, M.A. Basso

**Affiliations:** Fuster Laboratory of Cognitive Neuroscience, Department of Psychiatry and Biobehavioral Sciences, The Jane and Terry Semel Institute for Neuroscience and Human Behavior, David Geffen School of Medicine, UCLA Los Angeles, CA 90095 USA

## Abstract

In contrast to predictions from the current model of basal ganglia (BG) function, we report here that increasing inhibition from the BG to the superior colliculus (SC) through the substantia nigra (nigra) using *in vivo* optogenetic activation of GABAergic terminals in mice, produces contralateral orienting movements. Orienting movements resulting from activation of inhibitory nigral terminals are unexpected because decreases and not increases, in nigral activity are generally associated with orienting movements. To determine how orienting movements may result from activation of inhibitory terminals, we performed a series of slice experiments and found that the same optogenetic stimulation of nigral terminals used *in vivo*, evoked post-inhibitory rebound depolarization and spiking in SC output neurons *in vitro*. Only high frequency (100Hz) stimulation evoked contralateral movements *in vivo* and triggered rebound spiking *in vitro*. The latency of orienting movements relative to the stimulation *in vivo* was similar to the latency of rebound spiking *in vitro*. Taken together, our results point toward a novel hypothesis that inhibition from the BG may play an active rather than passive role in the generation of orienting movements in mice.

## INTRODUCTION

Stemming from experiments performed in monkeys, cats and rodents in the late 80’s and using traditional electrophysiological and anatomical approaches (Hikosaka and Wurtz, 1985a) a model of the role of the BG in movement generation in health and disease emerged (Albin et al., 1989; DeLong, 1983, 1990; Penney and Young, 1983). The model, currently described in all major neuroscience textbooks, proposes that the basal ganglia (BG) play a permissive role in the generation of movement through modulation of the amount of inhibition on BG target structures such as the thalamus and superior colliculus (SC). For orienting movements specifically, one of two output nuclei of the BG, the substantia nigra pars reticulata (nigra) contains GABAergic neurons that project primarily to the ipsilateral SC, a key structure involved in the control of orienting (Deniau et al., 2007; Liu and Basso, 2008; Sato and Hikosaka, 2002). Nigral neurons are active tonically with rates ∼50-100 spikes/sec and as such, provide a constant suppression of SC neuronal activity (Deniau et al., 2007). The removal of nigral inhibition on the SC, combined with an excitatory drive from the cerebral cortex to SC, produces contralateral orienting movements. The cascade of disinhibition to the SC is mediated by the direct BG pathway and is offset by activation of the indirect BG pathway, which acts to suppress movements by increasing the inhibitory output of the nigra on the SC (Hikosaka et al., 2000). The disinhibitory nature of BG function enjoys extensive experimental support in a variety of species including monkeys, cats, rodents and even lampreys (Grillner and Robertson, 2016; Joseph and Boussaoud, 1985; Reiner et al., 1998; Stephenson-Jones et al., 2012). Disinhibition as a mode of BG action, is often referred to as the rate model as it is the rate of spiking in the inhibitory output neurons of the BG onto target structures that determines whether or not a movement commences; high rates and more inhibition result in no movement whereas low rates and less inhibition, result in movement (Albin et al., 1989; Nelson and Kreitzer, 2014).

Testing models of BG function at the circuit level is now possible with optogenetics. Recent work in mice using causal manipulations of the direct and indirect pathways, suggests that the BG play a supportive rather than permissive role in reaching movements, and that the direct and indirect pathways of the striatum operate in concert to produce movement rather than antagonistically, wherein one disinhibits and the other inhibits movement (Klaus et al., 2019; Tecuapetla et al., 2016; Yttri and Dudman, 2016). We reasoned we could use optogenetics to provide a causal test of the disinhibition model of orienting at the level of the output, via the nigro-collicular circuit. The prediction from the disinhibition model is that increasing the activity of the nigro-collicular circuit unilaterally should suppress contralateral movements. Because unilateral inhibition would create an imbalance between the two SCs, it is also possible that ipsilateral movements may occur. A previously not considered possibility, is that activation of the nigral inhibitory circuit results in activation of the SC through a process of post-inhibitory rebound depolarization (RD) (Kim et al., 2017; Person and Perkel, 2005). To arbitrate between these three possibilities, we used AAV injections, to express Chronos, an opsin with fast kinetics (Klapoetke et al., 2014) in the nigral inhibitory terminals and activated them in the SC of mice with blue light. Surprisingly, we found that unilateral optogenetic activation of nigral afferents in the SC, evoked movements contralateral to the side of stimulation, rather than suppressed contralateral movements as predicted by a model of disinhibition. To understand the mechanism of this unexpected result, we performed SC slice experiments and found that the same optogenetic stimulation of nigral terminals used *in vivo*, evoked post-inhibitory RD and spiking in SC output neurons *in vitro* through activation of T-type Ca^++^ channels. These results suggest that the control of orienting movements by the inhibitory nigro-collicular circuit, may be active as well as passive, pointing toward a need to revisit our understanding of the role of BG in orienting movements.

## RESULTS

### Optogenetic activation of nigral terminals in the SC evokes contralateral orienting movements in mice

The idea that more inhibition from the BG produces less movement and less inhibition produces more movement enjoys considerable experimental support, much of which is circumstantial or correlational (Basso and Sommer, 2011; Freeze et al., 2013; Hikosaka and Wurtz, 1983b, 1983a; Hikosaka et al., 2000; Klaus et al., 2019; Nelson and Kreitzer, 2014; Schmidt et al., 2013). We reasoned we could use optogenetics to activate the nigral afferents in the SC to increase the inhibition on to the SC, providing a direct, causal test of the hypothesis that more inhibition suppresses contralateral orienting movements. Figure 1a shows three possible outcomes. Unilateral activation of nigral afferents should increase the suppression of ipsilateral SC neurons, preventing contralateral movements (Figure 1a_1_). Activation may also disinhibit the contralateral SC through inhibitory commissural connections, which might result in ipsilateral movements (Figure 1a_2_). A third, previously unconsidered possibility, is that activation of the nigro-collicular inhibitory circuit generates post-inhibitory rebound depolarization (RD) and spiking in SC output neurons, evoking contralateral movements (Figure 1a_3_). To arbitrate between these possibilities, we performed unilateral viral injections of AAV9-Syn-Chronos-GFP and AAV9-Flex-Chronos-GFP into the nigra of wild-type or GAD2-Cre mice, respectively. The latter mice were used to target the nigral GABAergic neurons specifically. Using 470nm light through an implanted fiber optic ferrule, we activated the nigral inhibitory terminals located in the SC while mice explored an open field environment (Supplemental Figure 1).

**Figure 1.**
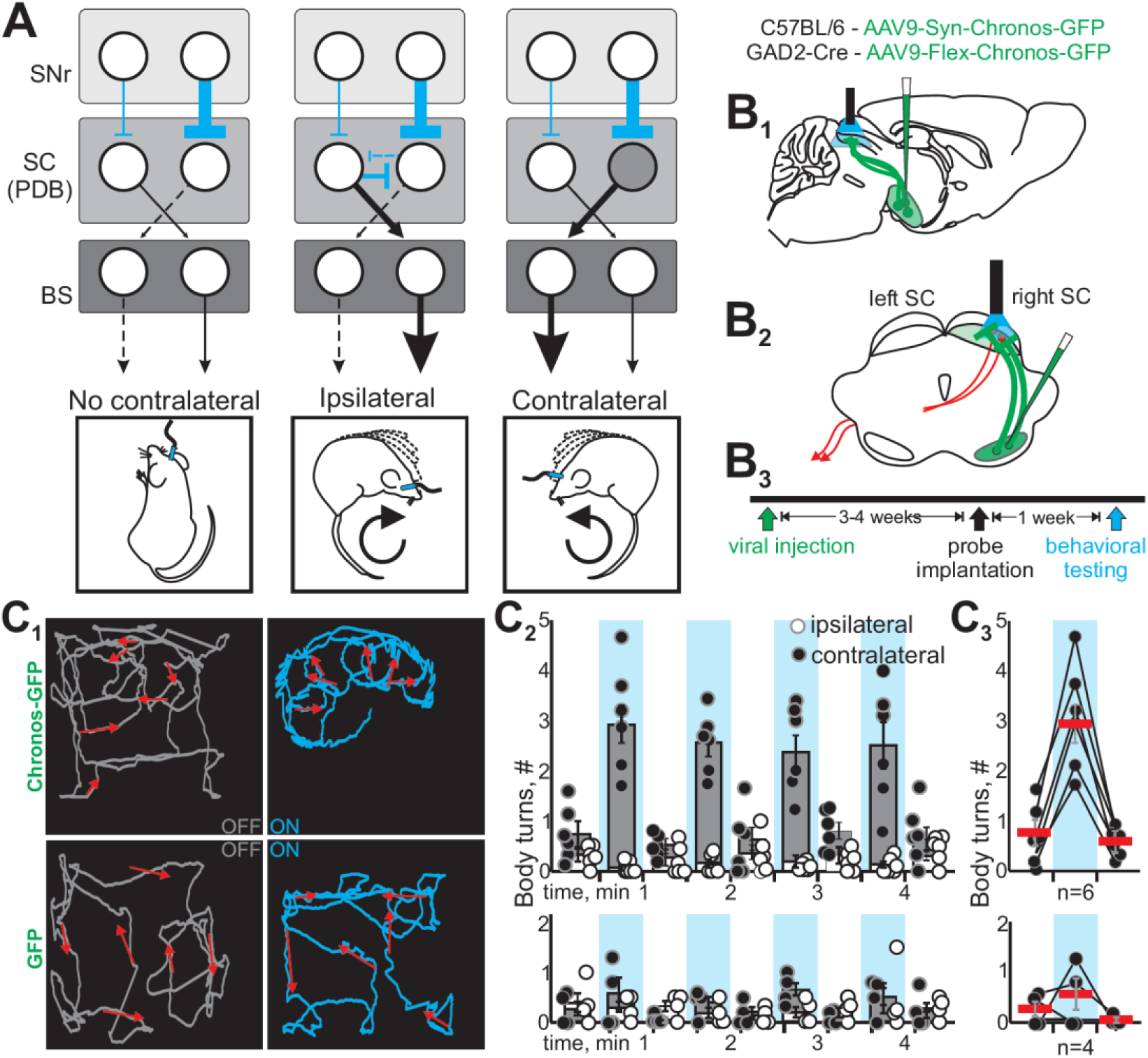
Optogenetic activation of the nigro-collicular circuit evokes contralateral orienting movements in mice. (**a**) Schematic depiction of the known synaptic connection between the nigra, the superior colliculus (SC) predorsal bundle neurons (PDB) and the brainstem and spinal cord orienting centers (BS) targeted by the PDB neurons of the SC. Cyan lines show inhibitory connections and black lines and arrowheads show excitatory connections. The thickness of the lines depicts the putative level of activity of the circuit with dashed lines indicating reduced activity. The circles represent the two nigrae, the two SCs and the two BS centers and the images below show predicted outcomes of optogenetic activation of the nigro-collicular circuit. (**a**_**1**_) The textbook model of nigro-collicular function - activation of the inhibitory nigro-collicular circuit inhibits the ipsilateral SC PDB neurons, preventing the generation of contralateral movements. (**a**_**2**_) A second possibility is that activation of the nigro-collicular circuit inhibits PDB neurons of the ipsilateral SC leading to disinhibition of the contralateral PDB neurons through commissural and interneuronal connections, resulting in a movement ipsilateral to the stimulation. (**a**_**3**_) A third possibility is that the cessation of the nigro-collicular activity produces spiking in the ipsilateral SC PDB neurons through rebound depolarization (RD, gray circle) causing contralateral movement. (**b**_**1**_) Schematics of a parasagittal section of a mouse brain showing the procedure for optogenetic activation of nigro-collicular terminals. Green shows the injection of either AAV9-Syn-Choronos-GFP or AAV9-Flex-Chronos-GFP into the nigra of wild type (C57BL/6) or GAD2-Cre mice, respectively. The blue light shows that we stimulated the Chronos expressing nigral terminals in the deep/intermediate SC on the same side as the injection of virus into the nigra. (**b**_**2**_) Schematic of a coronal section of a mouse brain showing the unilateral viral injection into the nigral and its terminals in the ipsilateral SC (green). Blue light shows the ipsilateral location relative to the nigral injection, of the probe and light stimulation. The red lines represent the contralateral projections of the collicular output neurons through the predorsal bundle. (**b**_**3**_) The timeline of injection and measurement for the experimental procedures. (**c**_**1**_) The top panels show movement traces, collapsed over time, depicting the tracked head position of a mouse injected with Chronos-GFP in an open field environment. The traces show the tracked interval of 30 seconds before (gray traces - OFF) and during (cyan traces - ON) optogenetic stimulation of nigral afferents in the SC. The bottom panels show the same measurement in a mouse injected with the control virus (GFP). The red arrows show the movement vector calculated on 5 seconds bouts of the corresponding tracking. The direction and length of each arrow depicts the direction and speed of movement for each interval. (**c**_**2**_) We quantified the number of ipsilateral (∘) and contralateral (●) turns relative to the stimulation site induced in mice. Gray and white bars (obscured by the white circles), show the average number of contra and ipsilateral body turns recorded in a 30s period during stimulation (100Hz, cyan shade) and during no stimulation (no shading) intervals for all the mice. Each point shows the mean number of turns for each interval measured over four sessions for each of the mice. The top panel shows the measurements made in Chronos-GFP mice (n=6) and the bottom panel shows the measurements for four control mice (n=4). (**c**_**3**_) The mean number of contralateral turns for all six mice (red horizontal bars, mean±SE) before, during (cyan shade) and after the first stimulation. Top panels show the quantification of body turns in Chronos-GFP mice and the bottom panels show the results from control mice.

Activation of the nigro-collicular terminals in mice expressing Chronos, unexpectedly evoked contralateral movements whereas mice injected with blank virus without Chronos, showed no discernible patterns of movement with stimulation (cf., Figure 1c_1_ top and bottom panels; Supplemental videos 1 and 2). The number of body turns contralateral and ipsilateral to the side of optogenetic stimulation recorded in the five OFF and four ON periods of a typical behavioral session is plotted for six mice expressing Chronos (Figure 1c_2_; top panels). During the stimulation-ON periods, the number of contralateral turns was significantly different from the number of ipsilateral turns (2×2 repeated measures ANOVA f(5,1)=58.73, p=0.001) whereas no differences were observed between contralateral and ipsilateral turns during the OFF periods (2×2 repeated measures ANOVA f(5,1)=4.01, p=0.102). No statistically significant differences were found for either the number of contralateral nor ipsilateral turns for control mice (Figure 1c_2_ bottom panel; 3×2 repeated measures ANOVA f(9,3)=0.09, p=0.962). The top panel in Figure 1c_3_ shows the number of contralateral turns for the first two OFF and the first ON periods for the six Chronos mice with their means (red bars; ON = 2.97±0.33; OFF = 0.78±0.23 and 0.54±0.092), indicating that all the mice injected with Chronos increased the number of contralateral turns upon light stimulation. The bottom panel of Figure 1c_3_ show the same for the four control mice (red bars: ON = 0.56±0.28; OFF = 0.21±0.13 and 0.05±0.023). In mice injected with Chronos, the mean number of ipsilateral turns for the first ON session was 0.03±0.03 whereas the means for the first and second OFF sessions were 0.22±0.108 and 0.19±0.11. Similar numbers were found for the control mice (ON = 0.21±0.084; OFF = 0.26±0.11 and 0.19±0.076). Thus, surprisingly, 100Hz optogenetic activation of the inhibitory nigro-collicular circuit in mice produces contralateral orienting movements rather than suppresses them as the model of disinhibition predicts.

### Predorsal bundle neurons show RD and RD-evoked spiking

It is unknown how activation of nigral inhibitory afferents in the SC could lead to a contralateral orienting movement. One possible hypothesis is that predorsal bundle (PDB) output neurons of the SC, are excited through post-inhibitory rebound depolarization (RD) evoked by GABA release from nigral terminals innervating the SC. Work in rodents and birds, indicates that pallidal (Area X) activation evokes RD and spiking in thalamic (DML) neurons (Kim et al., 2017; Person and Perkel, 2005), but see (Edgerton and Jaeger, 2014). We reasoned that a similar mechanism may be at play in the nigro-collicular circuit.

We first assessed whether PDB neurons were capable of producing RD and RD-evoked spiking by retrogradely labeling PDB neurons with injection of retroAAV2-CAG-tdTomato into the pontine reticular nucleus (PnC) and recording visually-identified PDB neurons using patch-clamp (Figure 2a_1-2_). 100pA positive current steps induced membrane depolarizations and triggered a train of action potentials in PDB neurons (Figure 2b_1_; black trace). 100pA step of negative current induced a sustained hyperpolarization at the end of which, the membrane voltage (V_m_) repolarized beyond its resting level and action potentials appeared (Figure 2b_1_; red trace). 85% (136/160) of PDB neurons showed RD-evoked spikes upon similar negative current steps, demonstrating that RD and RD-spikes induced by hyperpolarization are common in PDB neurons (Figure 2b_2_). To test whether RD in PDB neurons depended on the magnitude of the hyperpolarization, we recorded PDB neurons while injecting different magnitude current steps. Increasing the amplitude of negative current steps increased the amplitude of the hyperpolarization and evoked larger RDs in PDB neurons (Figure 2c_1_). To better observe the RD, we performed this experiment in the presence of 1μM TTX to block the RD-evoked spikes. The amplitude of the RD was largest in the condition with the largest negative current step (Figure 2c_1_; cf., black and blue traces and inset). All 6 PDB neurons tested showed a linear relationship between the amplitude of the step current and the RD amplitude (Figure 2c_2_; black and grey lines, mean R^2^ = 0.82 ± 0.064). We reasoned that if the RD induced by the negative step currents underlies the spikes seen at the end of the hyperpolarization, steps increasing the RD amplitude should increase the number of spikes. To test this, we performed the same experiment in the absence of TTX and found that the amplitude of the hyperpolarizing step also correlated with the number of RD-evoked spikes observed (Figure 2d_1_). The linear relationship for 6 PDB neurons appears in Figure 2d_2_ and the colored circles correspond to the traces in Figure 2d_1_ (mean R^2^ = 0.90 ± 0.028). These results demonstrate that in PDB neurons, both RD and RD-evoked spikes are common features and their amplitude and frequency are proportional to the magnitude of the preceding hyperpolarization consistent with reports from other neuronal cell types (Aizenman and Linden, 1999; Wang et al., 2016).

**Figure 2.**
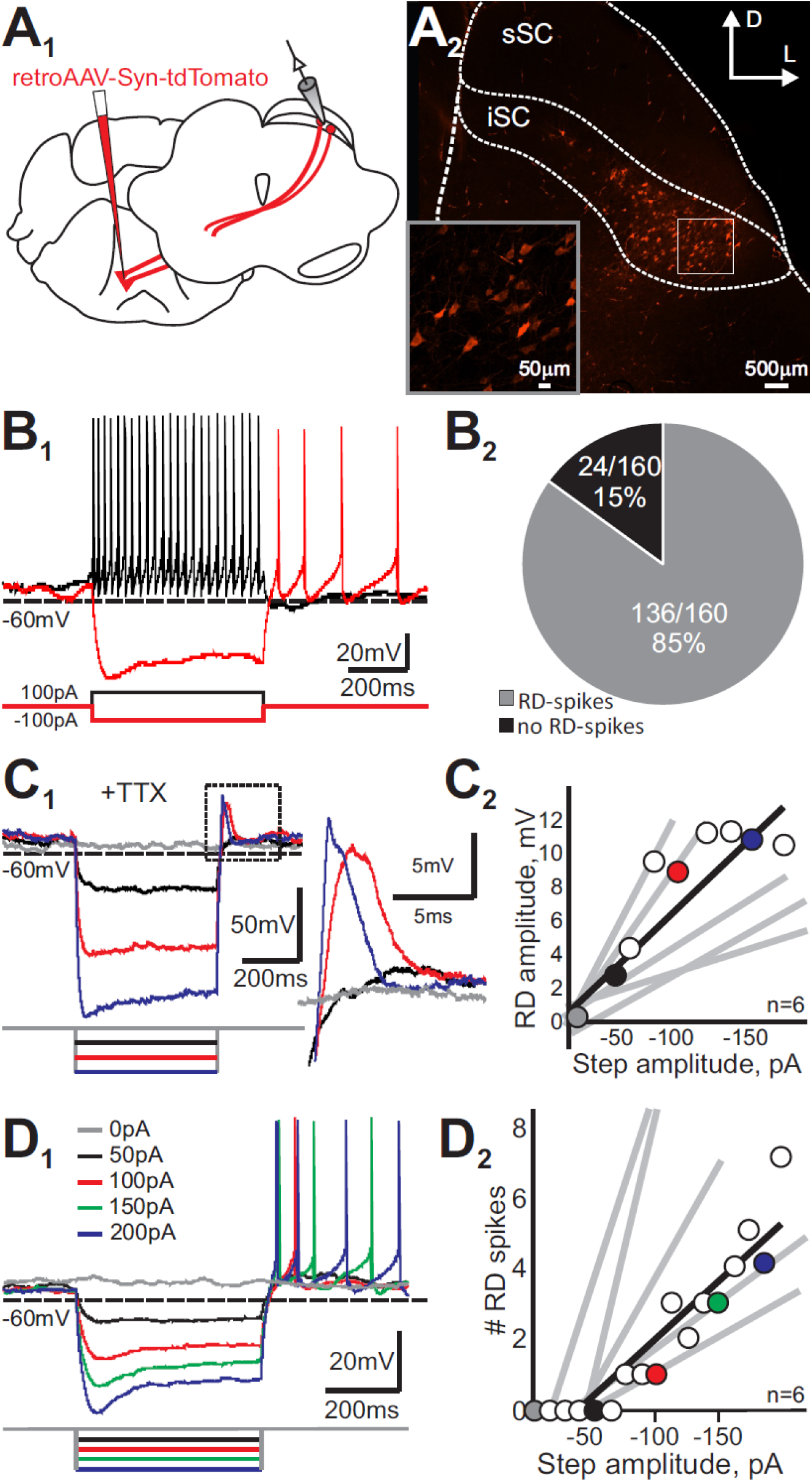
Predorsal bundle neurons (PDB) of the SC show robust RD. (**a**_**1**_) We injected 200-300nl of retroAAV-Syn-tdTomato into the caudal pons to label the contralateral PDB neurons of the SC retrogradely (red). (**a**_**2**_) Confocal image showing the distribution of PDB neurons in the SC. sSC = superficial SC; iSC = intermediate/deep SC. Scale bar, 500 μm. The inset shows a high magnification confocal image of the region of the iSC indicated by the box. Scale bar, 50μm. (**b**_**1**_) Current-clamp traces recorded from a PDB neuron. 500ms duration, +100pA current injections depolarized PDB neurons and evoked robust discharges of action potentials (black traces). 500ms, −100pA current injection induced sustained hyperpolarization followed by RD and spiking (red traces). (**b**_**2**_) Most PDB neurons exhibited RD-evoked spikes in response to 500ms duration, −100pA current injection. (**c**_**1**_) Current-clamp traces in response to varying amplitude, 500ms current steps (red −150pA, blue −100pA, black −50pA, gray 0pA) in the presence of 1μM TTX to eliminate the spikes and show only the RD. The part of the trace outlined by the dotted black square is magnified on the right to display the RD evoked by the negative currents. (**c**_**2**_) The peak amplitude of the RD in mV plotted against the amplitude of the negative step current in pA. Colored circles correspond to the colored traces shown in (**c**_**1**_). The black line shows the linear regression calculated from the data obtained from the neuron depicted in (**c**_**1**_). Gray lines show the linear regressions of five other PDB neurons. (**d**_**1**_) Current-clamp traces in response to varying amplitude, 500ms current steps in the absence of TTX to show the RD-evoked spikes in PDB neurons. (**d**_**2**_) The number of RD-evoked spikes plotted against the amplitude of the negative step current. Colored circles correspond to the colored traces shown in **d**_**1**_. The black line shows the linear regression calculated from the data obtained from the neuron depicted in (**d**_**1**_). Gray lines show the linear regression calculated for five other PDB neurons.

### Optogenetic activation of nigral inhibitory terminals in the SC induces RD and RD-evoked spiking in PDB neurons

We next sought to investigate whether inhibitory inputs from the nigra into the SC were capable of evoking similar patterns in PDB neurons as seen with current injection. We activated nigral terminals expressing Chronos opsin in the SC using light, while recording from visually-identified PDB neurons labeled with retro-AAV-Syn-tdTomato from a PnC injection (Figure 3a_1-3_; Supplemental Figure 2). Using patch-clamp recording, we found that activation of the nigro-collicular terminals with 10Hz light pulses in the SC produced reliable IPSPs in PDB neurons (Figure 3b_1_, red trace). Bath application of TTX to block Na^++^ channels and 4-AP to block K^+^ shunting demonstrated that the IPSPs induced by the activation of the nigro-collicular terminals in PDB neurons were monosynaptic (Figure 3b_1_, blue trace). Even though optogenetic activation of the nigro-collicular pathway induced reliable IPSPs in PDB neurons, 10Hz light stimulation pulses did not evoke RD nor spiking in PDB neurons reliably (Figure 3b_2_, red trace). However, 100Hz stimulation induced a large hyperpolarization, often accompanied by robust RDs (Figure 3b_3_, red trace and Supplemental Figure 3). In some PDB neurons the opto-induced hyperpolarization produced RD but did not produce spiking (11/29 PDB neurons). A possible explanation for the lack of spiking is that the light activation induced hyperpolarizations of too small amplitude. As shown before, the likelihood of producing RD-evoked spikes in PDB neurons was proportional to the amplitude of the preceding hyperpolarization. Thus, we reasoned that in neurons were light pulses were unable to produce RD-evoked spikes, increasing the amplitude of the opto-induced hyperpolarization should produce reliable RD-evoked spikes. One way to increase the amplitude is to enhance the driving force of the ions underlying the hyperpolarization by holding the neurons at slightly depolarized V_m_s. Injection of small, slow positive currents that induced subthreshold depolarizations together with 100Hz light pulse stimulation of nigral terminals, produced reliable trains of spikes in PDB neurons (Figure 3c_1_). Note that at more hyperpolarized V_m_s, the light pulses induced a reversal in the membrane voltage (Figure 3c_1_, black trace). The linear correlation between the amplitude of the hyperpolarization induced by light stimulation and the V_m_ demonstrated that the same intensity light pulses induced larger hyperpolarizations at more depolarized V_m_s (Figure 3c_2_; reversal potential = −67.79±0.88mV). Furthermore, as we observed with current injections, the number of spikes produced by light pulses was proportional to the amplitude of the hyperpolarization induced by optogenetic activation of the nigral terminals in all 13 PDB neurons tested (Figure 3c_3_). To ensure that the slow current used to depolarize the V_m_ was subthreshold and did not evoke action potentials before the light pulses, we performed longer ∼1.0 sec current-clamp recordings before the light pulses. As shown in Figure 3d, there were no action potentials before the light stimulation at the V_m_s where reliable RD-evoked spikes were obtained. These results demonstrate that PDB neurons receive monosynaptic inhibitory nigral inputs and 100Hz light activation of nigral terminals produces hyperpolarization followed by RD and RD-evoked action potentials. These findings point toward a possible new mechanism to explain how contralateral movements are evoked with inhibitory nigral activation as observed *in vivo*.

**Figure 3.**
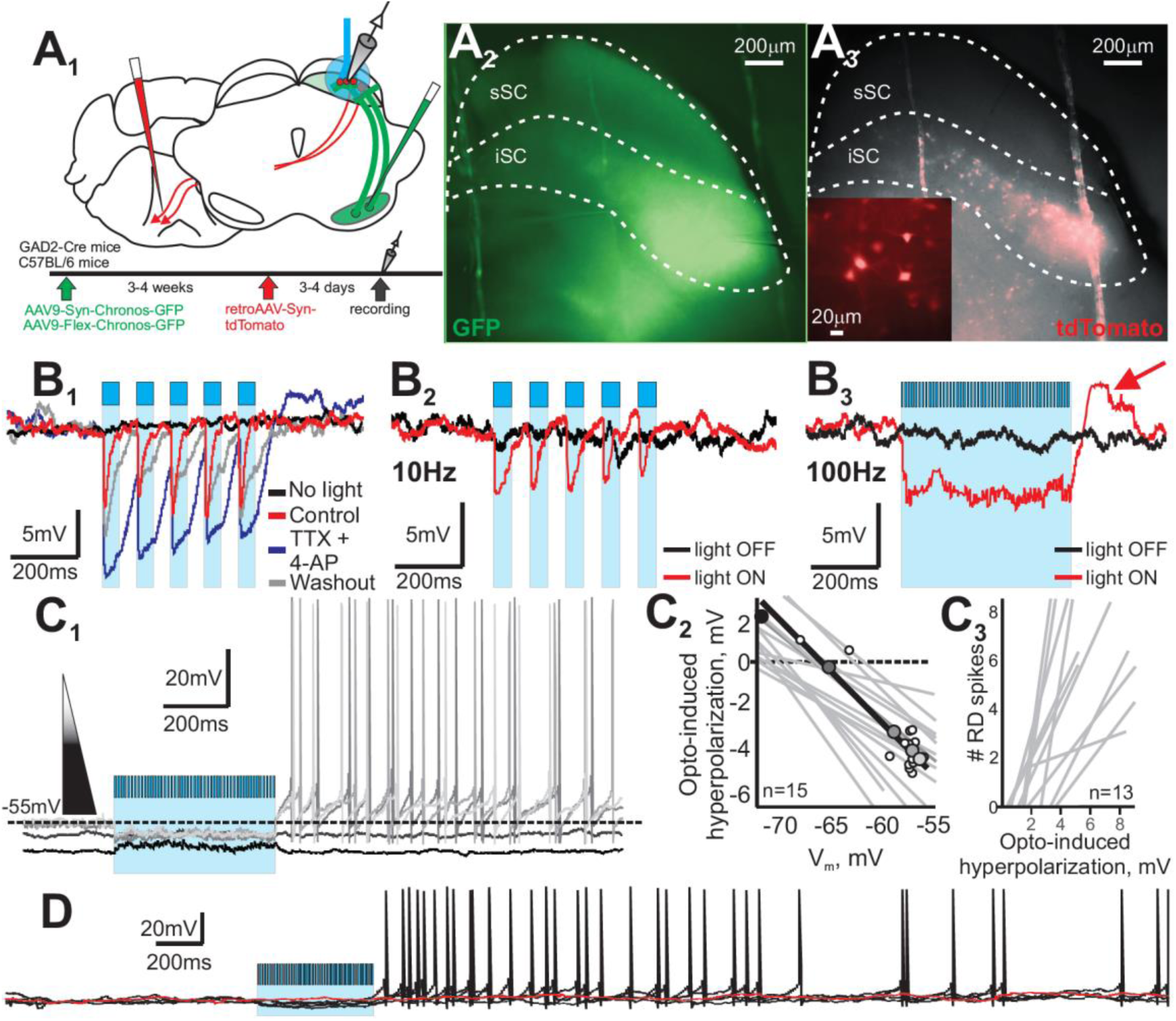
Optogenetic activation of GABAergic nigral inputs to the SC generates RD-evoked spikes in PDB neurons. (**a**_**1**_) Schematic coronal section showing the injection of either AAV9-Syn-Chronos-GFP or AAV9-Flex-Chronos-GFP into the nigra of CB57/BL or GAD2-Cre mice to transfect the nigro-collicular neurons and express Chronos in the nigral terminals located in the SC (green). Three to four weeks later, we injected retroAAV-Syn-tdTomato into the contralateral caudal pons to label PDB neurons ipsilateral to the nigra injections (red). (**a**_**2**_) Micrograph image of the fluorescing nigral fibers (GFP – pseudocolored green) in the iSC of a slice used to record from the retrogradely labeled PDB neurons. (**a**_**3**_) Micrograph image of the fluorescing PDB neurons (tdTomato - pseudocolored red) in same slice as **a**_**2**_. (**b**_**1**_) IPSPs from a PDB neuron (red trace) evoked by 10Hz light pulses (cyan bars). IPSPs remained after bath application of TTX/4-AP (black traces) and after 5-min of washout (gray traces), indicating that the IPSPs are monosynaptic. (**b**_**2**_) 10Hz light stimulation (cyan bars) evoked IPSPs in PDB neurons (red trace). (**b**_**3**_) In the same PDB neuron, 100Hz light stimulation (cyan shading) evoked a train of IPSPs (red trace) and a prominent RD at the end of the light stimulation (red arrow). Black traces are currents recorded without light stimulation. (**c**_**1**_) Five current-clamp traces recorded from a single PDB neuron at different V_m_s indicated by the shaded triangle. Darker traces indicate more hyperpolarized V_m_s. The cyan shading shows the time and duration of the 500ms duration, 100Hz optogenetic stimulation of the nigral terminals in the SC. Reliable RD and spiking appeared in PDB neurons when held at a slightly depolarized V_m_ (gray traces). (**c**_**2**_) The opto-induced hyperpolarization in mV is plotted against the V_m_. Shaded circles show the values obtained from the traces shown in **c**_**1**_. White circles show the rest of the data values recorded for the neuron shown in (**c**_**1**_) and the black line shows the calculated linear regression. Gray lines show the linear regressions obtained for 18 other PDB neurons. (**c**_**3**_) Number of RD-evoked spikes plotted against the amplitude of the opto-induced hyperpolarization in PDB neurons in mV. Gray lines show the linear regressions calculated for 13 PDB neurons. (**d**) Extended current clamp recording from a PDB neuron showing the current before (∼1.0 sec) and long after (∼3.0 sec) the optogenetic activation of nigral terminals. The black lines show four consecutive traces during light stimulation (cyan shading) and the red trace shows the V_m_ reference without light pulses.

### Features of RD and spiking in vitro predict features of orienting movements in vivo

100Hz optogenetic stimulation of inhibitory nigral terminals in the SC produced contralateral orienting movements in mice, and *in vitro*, produced post-inhibitory RD and RD-evoked spiking in PDB neurons of the SC. If post-inhibitory rebound spiking in PDB neurons is the physiological mechanism underlying the contralateral orienting movement seen *in vivo*, we reasoned that the features of the optogenetic activation resulting in spiking *in vitro* should be similar to those that produce orienting movements *in vivo*. We found that only high-frequency optogenetic stimulation of nigral terminals induced robust RD and RD-spikes (c.f. Figure 3b_2-3_ and Supplemental Figure 3a, b). 10Hz stimulation failed to evoke RD and spiking. Therefore, we predicted that 100Hz but not 10Hz unilateral optogenetic activation of nigral terminals *in vivo* would evoke movements contralateral to the site of stimulation. We recorded behavior in an open field apparatus during different stimulation patterns from four mice injected with Chronos into the nigra. (Supplemental Figure 1). Figure 4a depicts the number of contralateral movements triggered in periods with and without light stimulation with different stimulation patterns. As already described, 100Hz stimulation evoked consistent contralateral movement compared to intervals without light stimulation (2×2 repeated measure ANOVA f(3,1)=79.36, p=0.003). 10Hz or continuous light stimulation failed to evoke statistically significant contralateral orienting movements compared to intervals without stimulation (2×2 repeated measure ANOVA 10Hz: f(3,1)=0.67, p=0.474; continuous: f(3,1)=0.27, p=0.640). Moreover, none of these stimulation patterns evoked ipsilateral orienting movements (Figure 4b). Thus, only high frequency (100Hz) optogenetic stimulation of the nigral terminals in the SC, which evokes robust RD and spiking *in vitro*, evokes contralateral orienting movements *in vivo*. Light pulses of frequencies that induce small RD and no RD-evoked spikes *in vitro*, are ineffective at evoking orienting movements *in vivo*.

**Figure 4.**
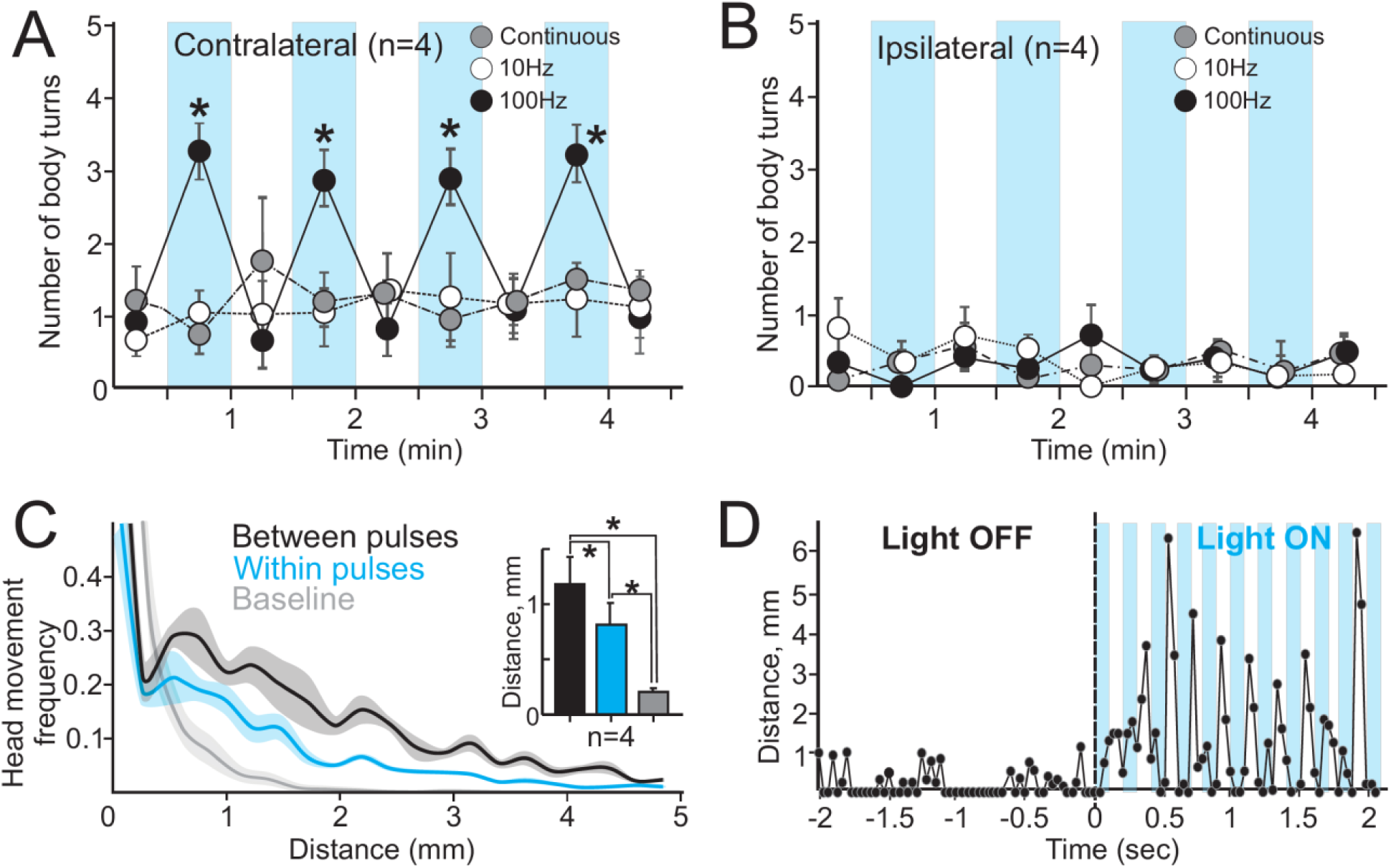
High frequency activation of GABAergic nigral terminals in the SC that evokes RD and spikes in PBD neurons *in vitro*, evokes contralateral orienting *in vivo*. (**a**) The number of body turns contralateral to the optogenetic stimulation of nigral terminals in the SC, in 30s intervals measured in an open field (see Methods) is plotted for different opto-stimulation frequencies (100Hz (●), 10Hz (∘) and continuous stimulation (●)). Cyan bars and white spaces represent the 30s intervals with and without opto-stimulation during the behavioral session, respectively. * = p<0.05 repeated measure ANOVA (3-way within-subject design), n = 4 mice. (**b**) Same as in (**a**) for ipsilateral body turns. (**c**) The normalized frequency of events (mouse head distance between two consecutive frames) and the value of the distance in millimeters for each head movement calculated during the first stimulation period. The lines show the averaged frequency of head movements calculated before the light stimulation (baseline, gray), within the light pulses (cyan) and between light pulses (black). The shaded areas show the ± SE (n=6). The inset shows the averaged distances for each of the conditions. * = paired t-test p<0.05. (**d**) Frame to frame distance in millimeters of the mouse head in a 4.0 sec video sample, during 100Hz optogenetic stimulation of nigral terminals. Data are plotted centered at the beginning of the light pulse stimulation. Cyan bars represent the 100ms light stimulation pulses and white spaces show the 100ms intervals between them. See also Supplemental Figure 1.

Another feature of the RD and RD-spikes observed *in vitro* is their timing - both appear at the end of the nigral activation and hyperpolarization of PDB neurons. If RD-evoked spiking in PDB neurons underlies the orienting movement after activation of the nigro-collicular pathway, the onset of the movement should correlate in time with the onset of the RD-evoked spiking in PDB neurons. We analyzed the head movements of the mice during the light stimulation to assess the timing. A slowed down video clip of two seconds before and after the initiation of 100Hz light stimulus shows that the initiation of the orienting movements with respect to the light stimulus occurred with a delay (Supplemental video 3). To confirm the delay in the initiation of movement with respect to the light, we measured the head position between consecutive video frames occurring during the light pulses and in between the light pulses for the first 30sec light stimulation (light ON). We then quantified the distance of the head positions between frames. If the initiation of the movement occurred in synch with the onset of the light pulses, most of the head movements should be seen in the video frames occurring during the light pulses. On the other hand, if there was a delay in the initiation of the movements, consistent with post-inhibitory rebound spiking, then most of the head movements should occur in the video frames between the light pulses. Figure 4c shows histograms of the number of head movements calculated between two consecutive video frames. Figure 4d shows the distance in millimeters the head traversed over time (see Methods). For the head movements recorded before the light stimulation, most traveled small distances (mean distance per head movement = 0.205±0.031 mm; Figure 4c, gray line). Within light pulses, we observed larger head movements (mean distance per head movement = 0.81±0.19 mm; Figure 4c, cyan line). However, the head movements recorded between the light pulses were larger compared to the those recorded within the light pulses (1.18±0.25 mm; paired t-test(5)=4.88, p=0.00455; Figure 4c, black line). Indeed, the largest head movements occurred between the light pulses (Figure 4d; Supplemental Figure 4). Figure 4d shows that the largest movements occurred within one or two frames after the cessation of the light pulse. Since the videos were recorded at ∼30.0 frames per sec, the occurrence of the movement at the end of the light pulse suggests the initiation of the head movements occurs within a 30-100ms window after the end of the light pulse.

We next measured the latency of the first RD-evoked spike in PBD neurons induced by either current injection or by optogenetic activation of the nigral terminals. In patch-clamp recordings, the first RD-evoked spike in 52 PDB neurons occurred with a median latency of 45.85ms and mean latency of 77.55±13.025ms after a step current injection. After optogenetic activation of nigral inhibitory terminals, the median latency of the first RD-evoked spike in 29 PBD neurons was 120.65ms and the mean was 112.79±12.86ms. The longer latency observed with optogenetic stimulation compared to the latency seen with current injection, is likely due to the smaller magnitude hyperpolarization that occurs with light. However, even with optogenetic activation, we detected a population of PDB neurons that showed RD-evoked spikes with a mean latency as short as of 44ms, similar to that recorded with current injections (Supplemental Figure 5). Taken together, we conclude that optogenetic activation of inhibitory nigral terminals in the SC results in contralateral orienting movements occurring between 30 and 100ms after stimulation, and that this time corresponds to the timing of post inhibitory RD-evoked spiking recorded *in vitro* consistent with the hypothesis that orienting movements evoked by nigral activation are mediated by post-inhibitory rebound spiking in PDB neurons of the SC.

### T-type Ca^++^ channels underlie the post-inhibitory RD in PDB neurons of the SC

Having shown that unilateral optogenetic activation of the nigro-collicular circuit generates contralateral orienting movements *in vivo*, likely due to post-inhibitory RD and spiking in PDB neurons, we next aimed to investigate the ionic mechanisms of the post-inhibitory RD and spiking in PDB neurons. One key possibility is that the hyperpolarization produced by nigral terminal stimulation, activates the non-specific cation channel, hyperpolarization-activated cyclic nucleotide-gated channel (HCN) leading to an influx of cations (I_h_). Activation of HCN in turn would lead to the activation of T-type Ca^++^ channels driving the V_m_ to potentials sufficient to activate Na^++^ channels and spiking.

We first assessed whether there was any evidence that PDB neurons present I_h_. In many neuronal types, a trademark of I_h_ is a voltage sag seen upon injection of negative current (Maccaferri and McBain, 1996; McCormick and Pape, 1990; Pape, 1996). We found that 89% (33/37) of wide field vertical (WFV) neurons of the superficial SC show the voltage sag, yet only 21.6% (8/37) of PDB neurons showed a voltage sag and the sag in PDB neurons was smaller than that seen in WFV neurons (Endo et al., 2008)(Supplemental Figure 6). In PDB neurons showing a voltage sag, bath application of the I_h_ blocker ZD 7288 (50μM) completely inhibited the sag (Figure 5a-b; mean sag amplitude = 3.32±0.46mV; mean sag amplitude + ZD 7288 = −0.17±0.66mV; n=7; paired t-test(6)=3.57; p=0.011), but failed to reduce the RD or the RD-evoked spiking occurring at the end of the negative current step (Figure 5a, c and d). The mean RD amplitude before and after ZD7288 application was 4.057±0.54mV and 4.54±0.80mV respectively (paired t-test(5) =-0.69; p=0.53). Thus, we conclude that I_h_ is unlikely to be a main contributor to post-inhibitory rebound spiking in PDB neurons of the SC.

**Figure 5.**
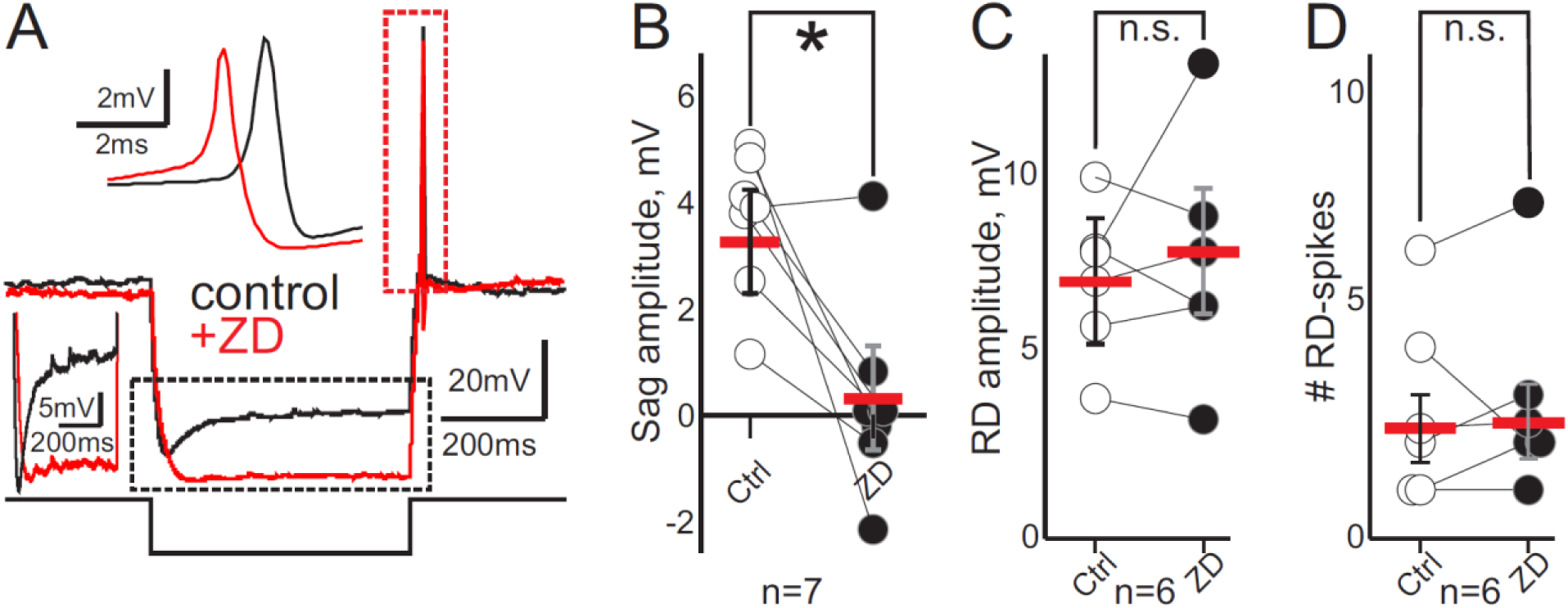
I_h_ does not contribute to the RD and RD-spikes in PDB neurons. (**a**) Current-clamp traces from a PDB neuron showing the changes in voltage upon a 500ms duration, −100pA negative current step (bottom black trace) before (control, black trace) and after (+ZD, red trace) bath application of the I_h_ blocker ZD 7288. The part of the trace outlined by the dotted black rectangle is compressed to highlight the difference in the sag (cf., black and red traces). The top inset expands the part of the traces outlined by the red dotted rectangle. (**b-c**) The sag (n=7) and RD amplitude (n=6) in mV during control (open circles) and during bath application of ZD (black circles) (n=7). Red bars, mean ± SE. *=p<0.05, paired t-test (**d**) Number of RD-spikes after negative step currents in PBD neurons in control and after ZD bath application (open and black circles, respectively). Red bars, mean ± SE. *=p<0.05, paired t-test.

We next asked whether T-type Ca^++^ channels play a role in producing post-inhibitory RD and spiking in PDB neurons as they appear to do in other neuronal cell types (Jahnsen and Llinás, 1984b, 1984a; Llinás and Jahnsen, 1982; Lo et al., 1998; Wang et al., 2016). After de-inactivation of the channels at hyperpolarizing voltages, T-type Ca^++^ currents appear when the membrane is depolarized above −60mV (Perez-Reyes, 2003). Thus, if T-type Ca^++^ channels underlie the appearance of RD and RD-evoked spiking in PDB neurons, both the RD and RD-spikes should be modulated by voltage in a manner similar to that seen for T-type Ca^++^ channel currents. We first recorded from PDB neurons at different V_m_s while applying a constant negative current step. Figure 6a shows that a 500ms duration, −100pA negative current step evoked a sustained hyperpolarization with a small RD when the PDB neuron was held at a hyperpolarized V_m_ (−67mV, Figure 6a bottom, black trace). As the membrane voltage became more depolarized, the same negative current step induced larger post-inhibitory RD and ultimately triggered RD-spikes (Figure 6a, lighter traces). We quantified the results of this experiment by measuring the hyperpolarization amplitude (Figure 6b, top left; R^2^=0.13±0.048), the latency of the first RD-evoked spike (Figure 6, top right; R^2^=0.69±0.12), the amplitude of the RD (Figure 6b, bottom left; R^2^=0.73±0.076) and the number of RD-evoked spikes (Figure 6b, bottom right; R^2^=0.66±0.096) at different V_m_s. All the R^2^ values were statistically different from zero except for the hyperpolarization amplitude (paired t-test(5)=2.51; p=0.054 hyperpolarization; paired t-test(5) = 6.22, p = 0.01 spike latency; paired t-test(5)=9.80, p=0.001 RD amplitude; t-test(5)=6.28, p=0.002 #RD-spikes). Although the amplitude of the hyperpolarization remained constant, depolarizing V_m_s increased the RD amplitude, reduced the latency of the first RD-evoked spike and increased the number of spikes elicited by the negative current step, consistent with increased activation of T-type Ca^++^ channels.

**Figure 6.**
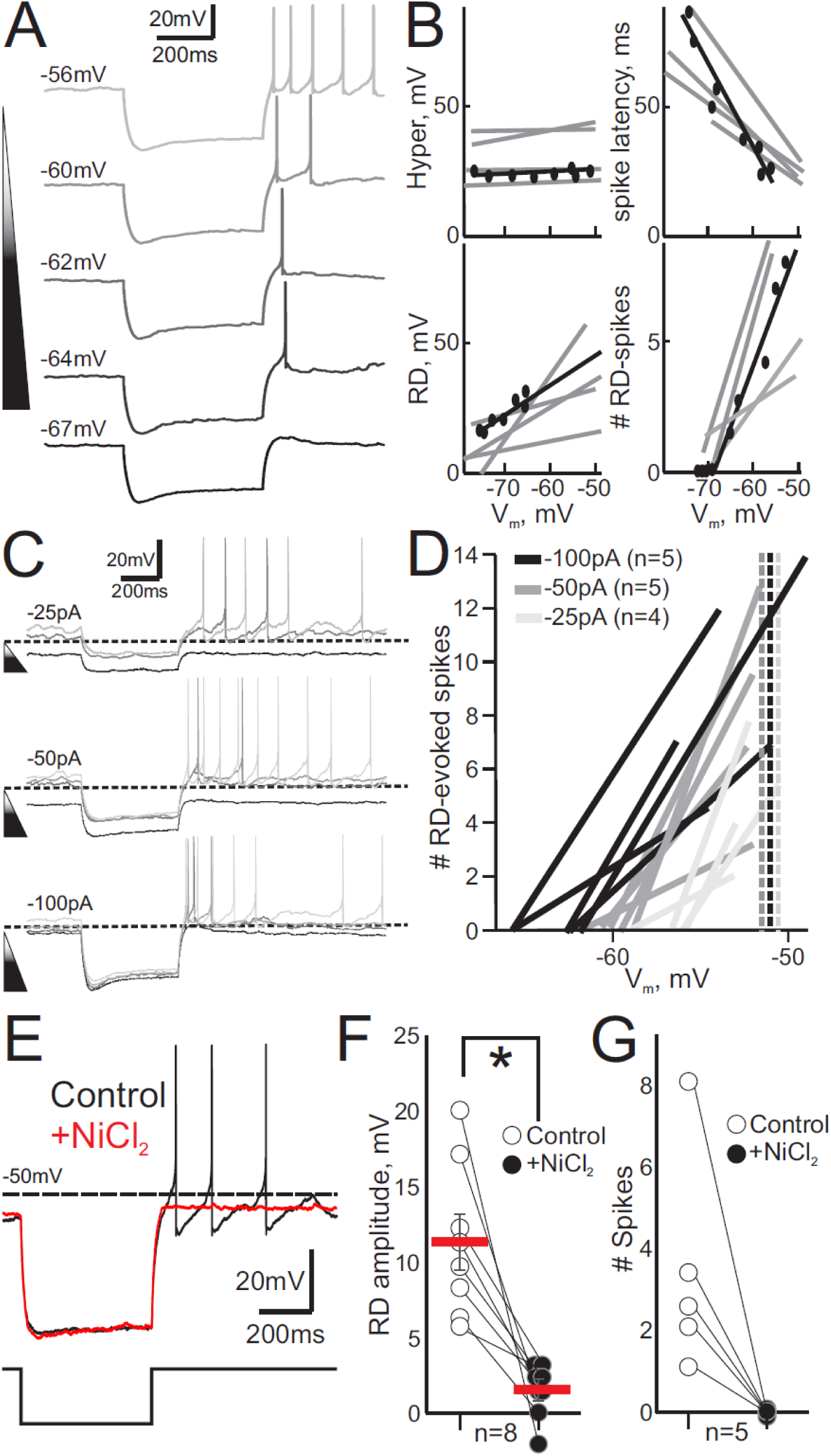
T-type Ca^++^ channels mediate RD and spiking in PDB neurons of the SC. (**a**) Current-clamp traces from a single PDB neuron plotted over time showing the voltage response to a 500ms −100pA negative current steps at different V_m_. The shaded triangle and traces indicate the V_m_ from hyperpolarized, −67mV (darker) to more depolarized −56mV (lighter). **(b)** Electrophysiological parameters induced by the negative current injection plotted against the V_m_. The black circles represent the data obtained from the PDB neuron shown in (**a**). The lines represent the linear regression calculated from those data. The gray lines show the parameters obtained for four other PDB neurons. The top left and right panels show the magnitude of the hyperpolarization induced by the negative current injection (Hyper, mV) and the latency of the first RD-spike in ms. The bottom left and right panels show the RD amplitude and the number of RD-evoked spikes. (**c**) Current-clamp traces plotted over time for a PDB neuron at different V_m_ in response to 500ms negative current steps of −25pA top traces, −50pA middle traces and −100pA bottom traces. Resting V_m_ = −57mV. Shaded traces represent V_m_ from hyperpolarized (darker) to depolarized (lighter). The dotted line is −60mV for calibration. (**d**) The number of RD-evoked spikes measured in a 1.0 sec epoch at the end of a current step plotted against the V_m_. The lines show the linear regression calculated from the number of RD-spikes recorded at each V_m_ evoked by each of the 500ms current steps (black −100pA (n=5), gray −50pA (n=5), light gray −25pA (n=4)). Dotted vertical lines show the average V_m_ for each of the current steps at which spontaneous spiking appeared. (**e**) Current clamp traces showing the changes in voltage after a negative 500ms, −100pA current step during control (black trace) and after application 500μm of the T-type Ca^++^ channel blocker NiCl_2_ (red trace). (**f**) RD amplitude induced by the injection of 500ms, −100pA negative current in PDB neurons in control (open circles) and during application of NiCl_2_ (black circles). Red bars, mean ± SE. (n=8). * = p<0.05, paired t-test. (**g**) The number of RD-evoked action potentials (spikes) after 500ms, −100pA negative current step injections in PDB neurons in control (open circles) and after bath application of NiCl_2_ (black circles; n=5).

Next, we recorded from PDB neurons at different V_m_s while applying negative current steps −25pA, −50pA and −100pA, to assess how V_m_ and hyperpolarization amplitude interact. Figure 6c shows a series of current-clamp traces recorded from a single PDB neuron. Applying a 500ms duration current step of −25pA, resulted in a small hyperpolarization and RD, and no RD-spikes (Figure 6c, top, black traces). The same current step at more depolarized V_m_s increased the RD amplitude and eventually generated RD-spikes (Figure 6c, top, grey traces). We found similar results with a −50pA current step (Figure 6c, middle traces cf., black and grey traces). A larger step current of - 100pA resulted in RD-evoked spikes at more hyperpolarized V_m_s, compared with the two previous step currents (Figure 6c, bottom traces). Figure 6d shows the linear regression calculated from the number of RD-evoked spikes at different V_m_ evoked by the hyperpolarizations induced by three different step currents. From these data, we calculated the V_m_ value at which the first RD-spike appeared for each of the different step current used; −100pA = −62.41±0.81mV; −50pA = - 58.69±0.39mV; −25pA = −54.67±2.052mV. These values demonstrate that larger hyperpolarizations increase the likelihood of evoking RD-evoked spikes at more hyperpolarized V_m_s. Conversely, smaller hyperpolarizations only evoked RD-spikes at more depolarizing V_m_s, within the activation voltage of T-type Ca^++^ channels. As a final test of whether T-type Ca^++^ channels underlie RD and RD-evoked spiking in PDB neurons, we recorded voltage responses from PDB neurons to 500ms duration, −100pA step current injections before and after bath application of 500μM NiCl_2_, which is known to block T-type Ca^++^ channels (Lee et al., 1999; Obejero-Paz et al., 2008). Figure 6e shows an example recording from a PDB neuron with and without NiCl_2_. The current injection induced a hyperpolarization followed by RD and RD-evoked spikes, both of which were suppressed by bath application of NiCl_2_ (Figure 6e, red traces; Supplemental Figure 7). The mean RD amplitude before and after NiCl_2_ were 11.30±1.90mV and 1.55±0.67mV respectively, and these differences were statistically significant (t-test(6) = 3.82; p = 0.0087). Bath application of NiCl_2_ also decreased the number of RD-evoked spikes from 3.6±1 spikes to 0 (t-test(4)= .98; p=0.020). Based on these combined findings, we conclude that post-inhibitory RD and RD-evoked spikes produced in PDB neurons by activation of the inhibitory nigral terminals in the SC result from de-inactivation and activation of T-type Ca^++^ channels.

## DISCUSSION

We used targeted optogenetics to activate the inhibitory afferents arising from the nigra and terminating in the ipsilateral SC of mice. Unexpectedly, we found that unilateral *in vivo* activation of the inhibitory nigral afferents in the SC produced contralateral orienting movements rather than suppressed them, as the model of BG function based on disinhibition predicts. *In vitro* experiments support the hypothesis that nigral afferents produce a strong hyperpolarization in PDB neurons followed by a post-inhibitory RD and spiking mediated by T-type Ca^++^ channels. High frequency stimulation (100Hz) but not low frequency stimulation (10Hz) resulted in orienting behavior and *in vitro* post inhibitory RD and spiking in PDB neurons. Moreover, the timing of the RD and spiking onsets relative to the light stimulus measured *in vitro*, matched closely the timing of the contralateral movements observed *in vivo*. Taken together, we propose that in addition to disinhibition from the nigro-collicular circuit leading to orienting movements, a second possible mechanism for producing orienting movements is a direct activation of PDB neurons via post-inhibitory RD and spiking.

### Relationship to previous studies

Our use of the GAD2-Cre mouse and the opsin Chronos allowed us to activate the nigral inhibitory projections in the SC specifically and with temporal precision, thus, driving the activity of nigral axonal projections at frequencies matching the *in vivo* discharge rates of the neurons. Direct activation of the circuit provides a causal test of the role of the nigro-collicular circuit in orienting movements. Other areas such as the GABAergic zona incerta (ZI) also project to the SC (Doykos et al., 2020; Hormigo et al., 2020; Kim et al., 1992; May and Basso, 2018; Watson et al., 2015). Our results, however, are unlikely attributable to the ZI projection as this is located ∼1.0mm anterior to our injection sites, and it lies ∼2mm dorsal to and at shallower angle than the nigra and our injection locations. Furthermore, despite the abundant inhibitory projections from the ZI to the SC, the projection strength to PDB neurons appears to be minimal (Doykos et al., 2020). The use of the GAD2-Cre mouse also ensured that the activation was specific to GABA containing neurons and not spurious activation of the adjacent dopaminergic neurons of the substantia nigra pars compacta. Finally, because we stimulated axons located within the SC, we can also rule out interpretations based on spread of virus to other midbrain neurons that do not project to the SC, as would be a concern if we stimulated nigral neuronal cell bodies directly.

To our surprise, we found that activation of the inhibitory nigro-collicular circuit led to movement generation rather than movement suppression. The mice moved freely in an open field during the light stimulation, so presumably nigral neurons would be variously active and pausing while the mice moved about. The introduction of the light would cause the release of GABA onto SC neurons, suppressing contralateral movements, and possibly resulting in an ipsilateral bias. This kind of behavior is seen with local injections of the GABA-A agonist muscimol into the SC (Hikosaka and Wurtz, 1983b, 1985b; Joseph and Boussaoud, 1985). In some ways, optogenetic stimulation of nigral afferents is a more biologically relevant test of the function of the nigro-collicular circuit than muscimol injections into the SC. First, optogenetic activation is a direct causal test of the role of the nigro-collicular circuit whereas muscimol injections provide circumstantial evidence because, as mentioned above, the SC receives inhibitory inputs from other brain areas in addition to the nigra, so muscimol effects could be as well explained by these other projections. Second, the SC contains inhibitory interneurons which would also be affected by muscimol injections but is avoided by targeted activation of nigral afferents. Third, muscimol is a potent GABA-A agonist with long-lasting effects, leaving neurons deeply hyperpolarized for hours (Edeline et al., 2002; Hikosaka and Wurtz, 1985a; Martin and Ghez, 1999). Optogenetic activation in contrast, allows for tighter temporal control of the stimulation and GABA release which, as our results showed, is needed for the triggering of RD and spiking in PDB neurons mediated by activation of T-Type Ca^++^ channels. The slow effect of muscimol injections in the SC would not provide the precise temporal control, maintaining the T-type Ca^++^ channels in PDB neurons in the inactivated state, preventing their de-inactivation and masking the occurrence of post-inhibitory RD and spiking in PDB neurons.

Post-inhibitory RD and spiking are found in many neuronal cell types including cerebellar nuclei neurons, thalamic neurons and even WFV neurons of the superficial SC (Chang et al., 2018; Endo et al., 2008; Hirano and Kawaguchi, 2014; Mitra and Miller, 2007; Person and Perkel, 2005; Sun and Wu, 2008; Wang et al., 2016). Electrical activation of the avian homologue of the mammalian pallidal nucleus, Area X, results in post-inhibitory RD and spiking in DLM neurons, the avian homologue of the mammalian motor thalamus (Person and Perkel, 2005, 2007). Follow up experiments, however, suggest that rebound spiking in the avian thalamus may stem from coincident excitation from cerebral cortex (Goldberg and Fee, 2012; Goldberg et al., 2012, 2013). We believe that coincident cortical activation is an unlikely explanation for the results of the nigro-collicular activation. In birds, the coincident activation observed using simultaneous neuronal recordings in Area X and DLM, appeared during bouts of singing (Goldberg and Fee, 2012; Goldberg et al., 2012). In our experiments, optogenetic activation of nigral afferents occurred at random times and thus was not correlated with any specific behavior in which coincident cortical drive would be associated. Moreover, many of the stimulation intervals occurred when the animal was at rest with minimal movement, presumably when cortical drives would also be minimal. Our results do not exclude the possibility that excitatory cortical inputs also trigger orienting movements during silencing of nigral inhibition. Our results do, however, provide strong evidence that nigral inhibition by itself is sufficient to drive spiking in SC PDB neurons and produce orienting. Thus, both disinhibition, and active post inhibitory rebound from the BG are likely to play a role in driving orienting movements.

Post-inhibitory RD and spiking dependent upon activation of T-type Ca^++^ appears in neurons of the mouse ventrolateral thalamus and is associated with neck and limb muscle contractions (Kim et al., 2017). The experiments performed in the mouse thalamus suggest that RD and spiking resulting from BG inhibitory input to the thalamus, play an active role in the control of movement and are not just associated with abnormal movement (Edgerton and Jaeger, 2014; Kim et al., 2017). Our results are consistent with this and show for the first time that the inhibitory input to the SC from the BG can play an active role in normal movement generation. Indeed, a role for nigral inhibition and post-inhibitory RD and spiking in PDB neurons of the SC helps explain a puzzling result in the monkey using electrical stimulation of the nigra (Basso and Liu, 2007; Liu and Basso, 2008). Electrical stimulation of the nigra in monkeys does not suppress visually-guided saccades but rather, results in a reduced and less variable saccadic reaction time, consistent with a synchronization of a large population of PBD neurons. Moreover, simultaneous recordings of SC neurons together with electrical activation of the nigra in monkey, reveal a transient pause in SC saccade-related neurons followed by a rapid rise in activity, again consistent with a hyperpolarization followed by RD and spiking, just as reported here (Liu and Basso, 2008).

### A new role for the nigro-collicular circuit in orienting behavior

The correlation between nigral pauses and SC bursts, together with findings from reversible inactivation studies, led to a model in which the pause of nigral neuronal activity served as a gate, allowing descending cortical excitatory inputs to the SC to drive the choice of a saccade (Sato and Hikosaka, 2002). Since those original experiments, recordings from nigral neurons, as well as pallidal neurons of the BG, reveal that the activity of BG output neurons is much more nuanced during saccades than the original gating model predicts (reviewed in (Basso and Sommer, 2011)). For example, some nigral neurons show pauses as soon as a visual stimulus appears and the pause in activity remains until the end of a saccade (Handel and Glimcher, 1999). Some nigral neurons show graded decreases in activity as the probability of a saccade changes (Basso and Wurtz, 2002). Still other nigral neurons show increases in activity around the time of a saccade (Handel and Glimcher, 1999). It is unclear how these varied response profiles from BG output neurons would fit into the traditional disinhibition model of orienting movement.

Recent work from the limb movement system is also changing the way we view the role of the inhibitory output of the BG. Rather than working in a push-pull antagonist framework, the increases and decreases in BG output nuclei activity are thought to work cooperatively to generate actions (Tecuapetla et al., 2016) and even to control the vigor of movement (Yttri and Dudman, 2016). Our results suggest that similar cooperative mechanisms may be at play in the orienting movement system controlled by the nigro-collicular circuit. One possibility is that the nigral neurons showing transient increases in activity work together with neurons that show pausing in activity. Coordinated action of these two neuronal types could lead to enhanced spiking and synchronization of a large population of SC neurons driving an orienting movement through rebound depolarization as well gating of excitatory inputs. A second not mutually exclusive possibility, is that the end of the pause in nigral neurons provides an important signal to SC neurons that also serves to synchronize the SC population activity to ensure a fast, coordinated orienting movement. A third possibility is that the pause in nigral activity combined with excitatory cortical (and/or cerebellar) inputs would generate subthreshold depolarizations that create a temporal window in which the likelihood of spiking is increased, similar to what we observed *in vitro* with changes in V_m_. The precise timing of the inhibition and excitation would provide a mechanism for enhancing signal to noise in the transformation of sensory signals to motor output.

## METHODS

### Animals and Surgery

All experimental protocols involving mice were approved by the UCLA Chancellor’s Animal Research Committee and complied with standards set by the Public Health Service policy on the humane care and use of laboratory animals, as well as all state and local guidelines. We used C57BL6 wild type mice (The Jackson Laboratory, stock no. 000664) and in GAD2-IRES-Cre knock-in mice (The Jackson Laboratory, stock no. 010802). 42 mice were used in the slice experiments and 12 mice were used in the behavioral experiments and all mice were maintained on a reverse circadian 12-h light/12-h dark cycle with food and water provided *ad libitum*. All surgical procedures were performed using aseptic technique and were performed using general anesthesia. Post-surgical analgesia was provided, and animal recovered from surgery for at least 5 days before experiments commenced.

### Statistical analysis

All the data are expressed as means and standard errors of the mean (SEM). We assessed the statistical significance using parametric tests when the data passed tests for normality (Kolmogorov-Smirnov test of normality) and significance was considered for test statistics with p value < 0.05.

See **ONLINE METHODS** for further details.

## ONLINE METHODS

### AAV vectors

For optogenetic activation of the nigral terminals in the SC (nigro-collicular), AAV9-Syn-Chronos-GFP (UNC, AV6102C), AAV9-Flex-Chronos-GFP (UNC, AV65553) were injected into the nigra of C57BL6 wild type or GAD2 mice. We used AAV9-Syn-EGFP (Addgene, 50465-AAV9) as expression control (blank virus). For retrograde labeling of PDB neurons, we used AAV2-retro-CAG-tdTomato (UNC, AV7643B).

### Stereotaxic injections

Mice were anesthetized with isoflurane (5.0% induction, 1.5-2.0% maintenance) and placed in a stereotaxic apparatus (Kopf Instruments, CA USA). A midline incision exposed the skull and the periosteum was removed. The skull was leveled using a digital display console (Kopf Instruments, CA, USA). Bregma and lambda were placed at the same z-position to ensure leveling in the pitch axis and the points AP:-3.20 and ML:+1.5/-1.5mm were placed at the same z-position to ensure leveling in the roll axis. We made a burr hole in the skull using a dental drill with a 0.5mm bit at the following coordinates AP:-3.4; ML: +3.2; DV:-4.3. Once the burr hole was done, the virus was loaded into a thin glass pipette connected to a 2.5 μl Hamilton syringe with a compression fitting set (cat# 55750-01, Hamilton, NV USA) and primed with mineral oil (VWR, cat# 9706-128). Aliquots of virus (2.0μl) were stored at −80°C and thawed at room temperature right before loading into the pipette. The injecting pipette was lowered to the calculated depth. Once in the target location, 200-300nl of virus was injected at a speed of 20-30nl/min using a QSI Stoelting microinjector (cat# 53311, Stoelting Co, IL, USA). The injection was performed at a 30° angle from vertical in the sagittal plane to reach the nigra and avoid the SC. The pipette remained at depth for 10 min following the injection before slowly retracting the pipette. After the injection, the skin was sutured and supplemental fluid as well as the analgesic carprofen were provided. The mice recovered in a controlled temperature recovery cage and were then returned to their home cage.

For the slice recording experiments, mice received a viral injection into the nigra as described above and then a second injection of retroAAV2-CAG-tdTomato into the caudal pons (PnC) 3-4 weeks later to label retrogradely the output neurons of the SC. The coordinates used for the PnC injections were, AP:-5.45; ML:-1.3; DV:-4.35. A 10° angle from vertical in the sagittal plane was used for the injection. Animals recovery and post-surgical treatments were completed as described above. Three to four days after the PnC injections, mice were anesthetized with isoflurane, euthanized and the brains were extracted for patch-clamp slice electrophysiological recording experiments as described below. To identify wide-field vertical (WFV) neurons of the superficial SC, we injected cholera-toxin b conjugated with Alexa Fluor 555 (ThermoFisher, cat #: C34776) into the pulvinar (Bickford et al., 2015). Visually identified WFV neurons in the ipsilateral SC labeled retrogradely were recorded using patch-clamp.

### Slice Electrophysiology

Preparation of acute brain slices was performed according to previous work (Villalobos et al., 2018). Brain slices containing the SC were prepared from adult mice (P30-P50) anesthetized with isoflurane before decapitation. The brains were quickly removed and cooled (4°C) in high sucrose cutting solution (in mM: 240 sucrose, 7 D-glucose, 7 MgCl_2_, 1.25 NaH_2_PO_4_, 2.5 KCl, 25 NaHCO_3_, 0.5 CaCl_2_ bubbled to saturation with 95% O_2_ – 5% CO_2_). Coronal brain slices (250-300μm) were cut using a vibratome (Leica VT 1200S) and subsequently transferred to a recovery chamber containing artificial cerebrospinal fluid (ACSF: 126 NaCl, 2.5 KCl, 26 NaHCO3, 1.25 NaH2PO4, 2 CaCl2, 2 MgCl2 and 10 glucose; ∼305 mOsm, pH 7.4) at 35°C for at least an hour before recording. Slices were then transferred to a submerged recording chamber on the stage of an upright Zeiss (Axio Examiner D1) microscope. Slices were superfused (2–3 ml/min) with standard ACSF and maintained at ∼30°C using an in-line heater controller (TC-324C, Warner Instruments). Individual PBD neurons were identified under infrared differential interference contrast (IR-DIC) and fluorescence using a Hamamatsu Cooled CCD camera (C11440-42U). Recordings were obtained using 3–5 MΩ electrodes filled with intracellular solution (K-Gluconate, 20 KCl, 0.2 EGTA, 2 MgCl_2_, 10 HEPES, 2 Na-ATP, and 0.5 Na-GTP; 290 mOsm; pH 7.3). Signals were amplified using a Multiclamp 700B amplifier (Molecular Devices, San Jose, CA, United States), digitized and stored on a PC. Series resistance and whole-cell capacitance were automatically compensated. To record PnC neurons at different V_m_s we applied slow constant current (∼5.0 sec) through the recording electrode. Electrophysiological data were analyzed using the Clampfit 10 software. To activate the nigro-collicular circuit optogenetically, we obtained slices from mice receiving nigral injections with Chronos-GFP. We used Chronos to activate nigral terminals, due to its fast kinetics (Klapoetke et al., 2014) and our desire to more closely match the *in vivo* discharge rates of nigral neurons which range from 50-125 spikes/sec (reviewed in (Basso and Sommer, 2011)). We illuminated the SC in slices through the 40X objective with a 470 nm LED light (∼5mW light output, Mightex). To produce light-evoked postsynaptic potentials in PDB neurons we presented light pulses at different frequencies, 100Hz and 10Hz using 500ms train duration. The pulse duration for the 100Hz stimulation was 5ms and for the 10Hz stimulation was 50ms. Similarly, as with the negative current steps, we assessed the RD amplitude and presence of RD-spikes at the end of the stimulation train. We also presented a single light pulse for a duration of 500ms in slices to match the continuous stimulation condition used *in vivo*, and this produced a hyperpolarization that decayed quickly over time, likely due to the activation kinetics of the opsin (data not shown). All chemicals were purchased from Sigma-Aldrich. To confirm the monosynaptic nature of the PSPs, recordings were obtained in the presence of 4-aminopyridine (4-AP, 20μM) and tetrodotoxin (TTX, 1μM). The sag amplitude of membrane voltage responses to negative step currents were measured as in (Tateno and Robinson, 2011). To test the ionic mechanism of the sag and the RD, we superfused ZD 7288 (ZD, 50μM) and NiCl_2_ (500μM) with ACSF. All recorded neurons were deemed healthy by assessing V_m_ magnitude and stability, and maximum spike voltage (e.g., < −50 mV and >0 mV, respectively).

### Optical Fiber Implantation

Three to four weeks after the nigra AAV injections, a ceramic ferrule with an optical fiber (0.5mm in diameter, 3.0mm length) was implanted with the fiber tip on top of the lateral portion of the intermediate/deep SC using stereotaxic coordinates (AP:-3.8, ML:+1.6, DV-2.2), ipsilateral to the side of the nigral injection. After five minutes to allow the probe to settle, the probe was affixed to the skull with consecutives layers of dental acrylic. Once the acrylic dried, the incision was closed with silk sutures. Recovery and post-surgical care were the same as described above. Behavioral testing commenced five - seven days after the optical fiber implantation. The right SC was implanted in 5 mice and the left SC was implanted in 1 mouse.

### Behavioral Measurement

Mice were handled daily by the experimenter for at least two - three days before making behavioral measurements. On the day of the behavioral measurement, mice were transferred to a room and were habituated to the room conditions for 15-20 minutes before the experiment started. Before each session, the apparatus was cleaned with 70% ethanol to eliminate odor cues. All behavioral measurements were made during the same circadian period (13:00-17:00) and were performed at the Behavioral Testing Core at UCLA (BTC-UCLA). Mice were habituated to the open field environment for 10-15 minutes before the start of each trial. The open field environment consisted of a transparent acrylic box (30 × 30 × 18 cm) and a zenithal video camera (DMK 22AUC03 ImagingSource, NC, USA) with a varifocal lens (Stoelting, UK), connected to a computer. The light pulses for optogenetic stimulation were controlled by an Ami-2 digital interface (Stoelting, UK) and powered by a Dual-Optogenetics-LED (Prizmatix, Israel) which provided a ∼25mW blue light output power. For every mouse, a behavioral session comprised of a five-minute recording consistent of 30s of baseline recording with no stimulation followed by consecutive 30s bouts of stimulation and no-stimulation intervals, repeated four times in a full session. In each set of experiments, we aimed to test the ability of varying frequencies of stimulation light to induce orienting movements *in vivo*. The stimulation protocols were selected based on the results of the *in vitro* stimulation and consist of a series of light ON/light OFF transitions, which correspond to the cessation of the inhibitory stimulus, where post-inhibitory spikes would be most likely to occur. Furthermore, these stimulation protocols allow us to rule out any other type of excitatory stimulation that could evoke orienting movements in our experiments, which would be reflected in movements induced by light stimulation regardless of the frequency. We presented light pulses at 100Hz, 10Hz and continuously. The pulse duration for the 100Hz and 10Hz stimulation were 5ms and 50ms, respectively (Supplemental Figure 1b_1-2_). For the 100Hz, the stimulation trains were presented in 100ms intervals, so in each 100ms interval there were ten light pulses with 5ms duration. Each 100ms stimulation interval was followed by 100ms with no stimulation, thereby producing ∼150 light ON/light OFF transitions during a single 30 sec bout of stimulation. For the 10Hz stimulation protocol, we used 1sec stimulation trains intervals, where each interval consisted of ten 50ms duration light pulses. In the 1sec interval there were the same amount of pulses as with the 100Hz protocol but at a lower frequency. Each 1sec stimulation interval was followed by a 1sec duration light OFF interval, producing 15 light ON/light OFF transitions during the 30sec bout of stimulation. The continuous stimulation protocol consisted of a single 30s light ON period. This protocol produced light ON/light OFF transitions only at the end of each 30s light stimulation (Supplemental Figure 1b_3_). The 30sec stimulation bouts were repeated 4 times during the testing sessions for all the stimulation protocols. The order of the stimulation protocol was randomized across all sessions. For each mouse, we averaged the number of body turns recorded in 30s bouts (light ON/light OFF) from four different sessions. For the head movements analysis, we averaged the values obtained from the histograms from 4 different mouse before, and during the first stimulation interval on a single behavioral session. At the end of each session, mice were returned to the housing cage and the open field was thoroughly cleaned with 70% ethanol before the next testing session. All behavioral data was recorded in the open field with the zenithal video camera (30 frames/sec) which monitored the instantaneous position of the mice. With the video taken by this camera the instantaneous position of the mouse, head, and tail, whole body turns, speed and time of light pulses were determined automatically using the AnyMaze software (Stoelting Co, IL, USA). For accuracy, we obtained from the videos the frame-by-frame absolute position of the head off-line using an in-house MATLAB program (code available upon request). The distance traveled by the mouse’s head between consecutive video frames (d) was calculated using the following formula: 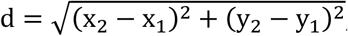, where x_1_ and y_1_ represent the horizontal and vertical position of the head in the open field during a set frame and x_2_ and y_2_ represent the horizontal and vertical position of the head recorded during the following video frame. Each coordinate value was aligned to the times of light ON or light OFF.

### Histological procedures

Mice were anesthetized with isoflurane and perfused with phosphate buffered saline (PBS) and PBS containing 4% paraformaldehyde (PFA). Brains were removed and incubated in PFA overnight. After rinsing with PBS, the brains were cryoprotected in 30% sucrose in PBS until sinking. Brains were then embedded in Tissue-Plus O.C.T. Compound (cat# 23-730-571, Fisher Scientific), frozen on dry ice, and cryo-sectioned coronally. 30-60μm thick sections were mounted on slides and cover-slipped with Fluoro-Gel mounting medium with TES buffer (EMS, 17985-30). Sections were scanned with an LSM 800 (Carl Zeiss) for confocal images or a Leica DMI8 (Leica Microsystems).

## Acknowledgements

We are grateful to Drs. Martha Bickford, Craig Evinger and Paul May for critical comments on the manuscript. We are grateful to Dr. Martha Bickford for assistance with the initial optogenetic experiments and Psyche Lee for assistance with the initial injections, Eduardo Alvarez and McKenna Lah for animal care, Kelly Zhang for technical support and Vaibhav Thakur for statistical assistance. We are also grateful to Irina Zhuravka, Dr. Lindsay Lueptow and the Behavioral Training Core at UCLA for help with the *in vivo* experiments. This work was supported by a Marion Bowen neurobiology postdoctoral fellowship grant to CAV and EY019663 and EY024153 to MAB.

## SUPPLEMENTAL MATERIALS

**Supplemental Figure 1.**
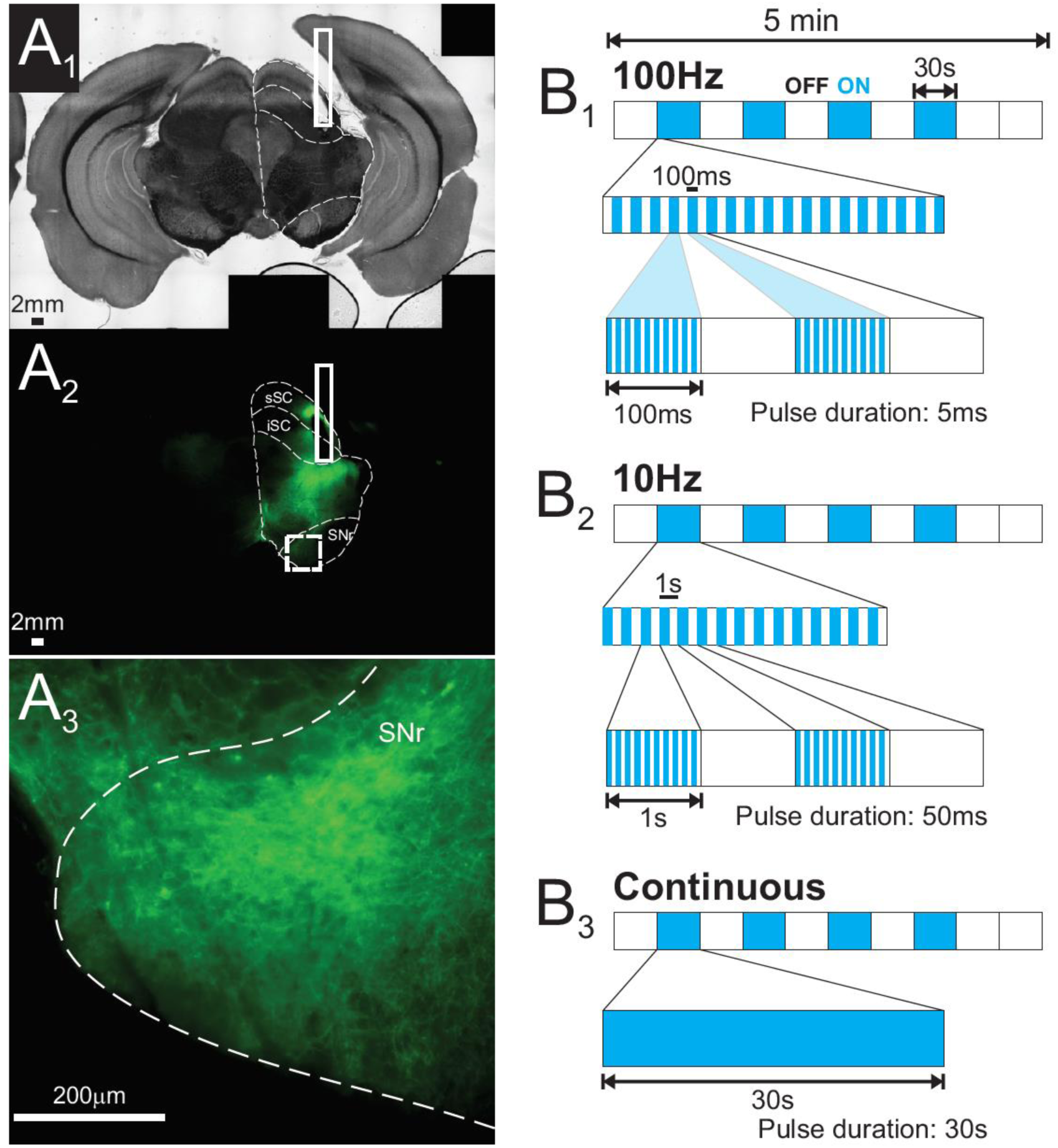
(**a**_**1**_) A bright field photomicrograph of a coronal section of a brain slice obtained from a mouse after performing the behavioral measurement and optogenetic stimulation. The solid white rectangle indicates the location of the optic fiber above the lateral iSC. Black bar = 2mm. (**a**_**2**_) Fluorescent micrograph (GFP) of the same slice shown in (**a**). As in (**a**), the solid rectangle shows the location of the optic fiber. Note that this area is not green because the fiber optic probe is there. The dashed square represents the area of the nigra amplified in (**a**_**3**_). (**a**_**3**_) Fluorescent micrograph (20X - GFP) of the section marked by the dashed square in (**a**_**2**_). White bar = 200µm. Referring to Figure 1 of the main text. (**b**) Schematics representing the different patterns of light stimulation used during the behavioral sessions. A single 5 min session consists of a 30 sec baseline of no stimulation, followed by consecutive 30 sec periods of stimulation (light ON, cyan squares) and no stimulation (light OFF, white squares) at different frequencies. The different stimulation patterns were aimed to increase or decrease the amount of light ON/OFF transitions, likely where the RD is triggered, without changing the overall length of stimulation (**b**_**1**_) For the experiments with 100Hz stimulation, during the ON period, the stimulation pattern consisted of a series of 100ms intervals with and without light. Within each interval with light pulses, a sequence of 100Hz light pulses each of 10ms duration, were triggered. During the OFF periods, there was no stimulation. (**b**_**2**_) For the experiments with 10Hz stimulation, during the ON periods the pattern consisted of a series of 100ms intervals with and without light. Within each interval with light pulses, a sequence of 10Hz light pulses each of 100ms duration, were triggered. (**b**_**3**_) For the experiments with continuous stimulation, the light pulse equaled the length of the ON period of 30 sec.

**Supplemental Figure 2.**
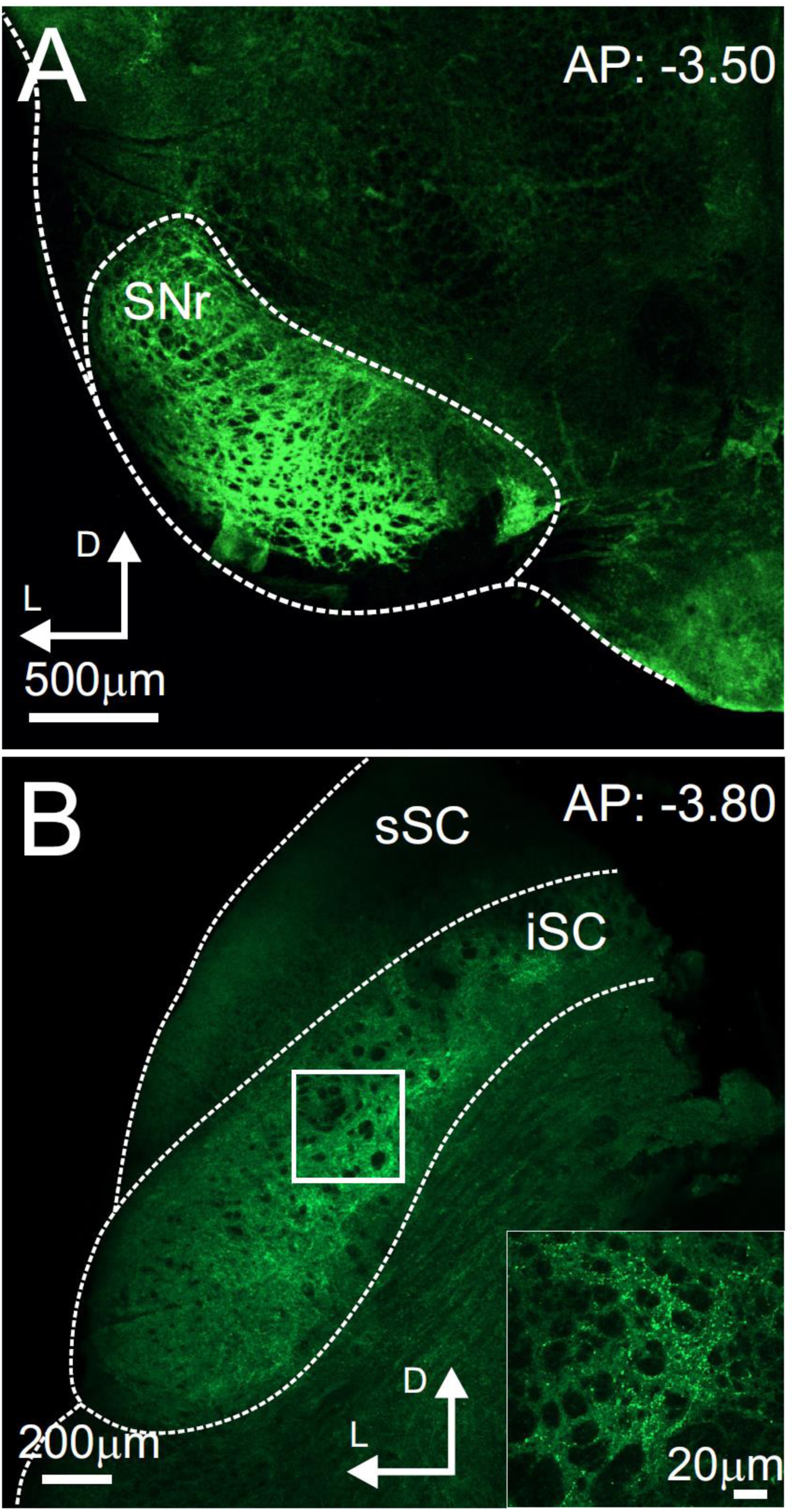
Distribution of the nigral terminals in the SC. (a) Confocal fluorescent micrography of a coronal slice (AP: −3.50mm from bregma) showing the expression of the Chronos-GFP virus in the nigra. Scale bar is 500µm. (**b**) Confocal fluorescent micrograph of a coronal slice (AP: −3.80mm from bregma) showing the distribution of nigral terminals in the iSC. Scale bar is 200µm. The solid square shows the region within the iSC where the 40x high magnification image in the inset was obtained. Scale bar is 20µm. Referring to Figure 4 of the main text.

**Supplemental Figure 3.**
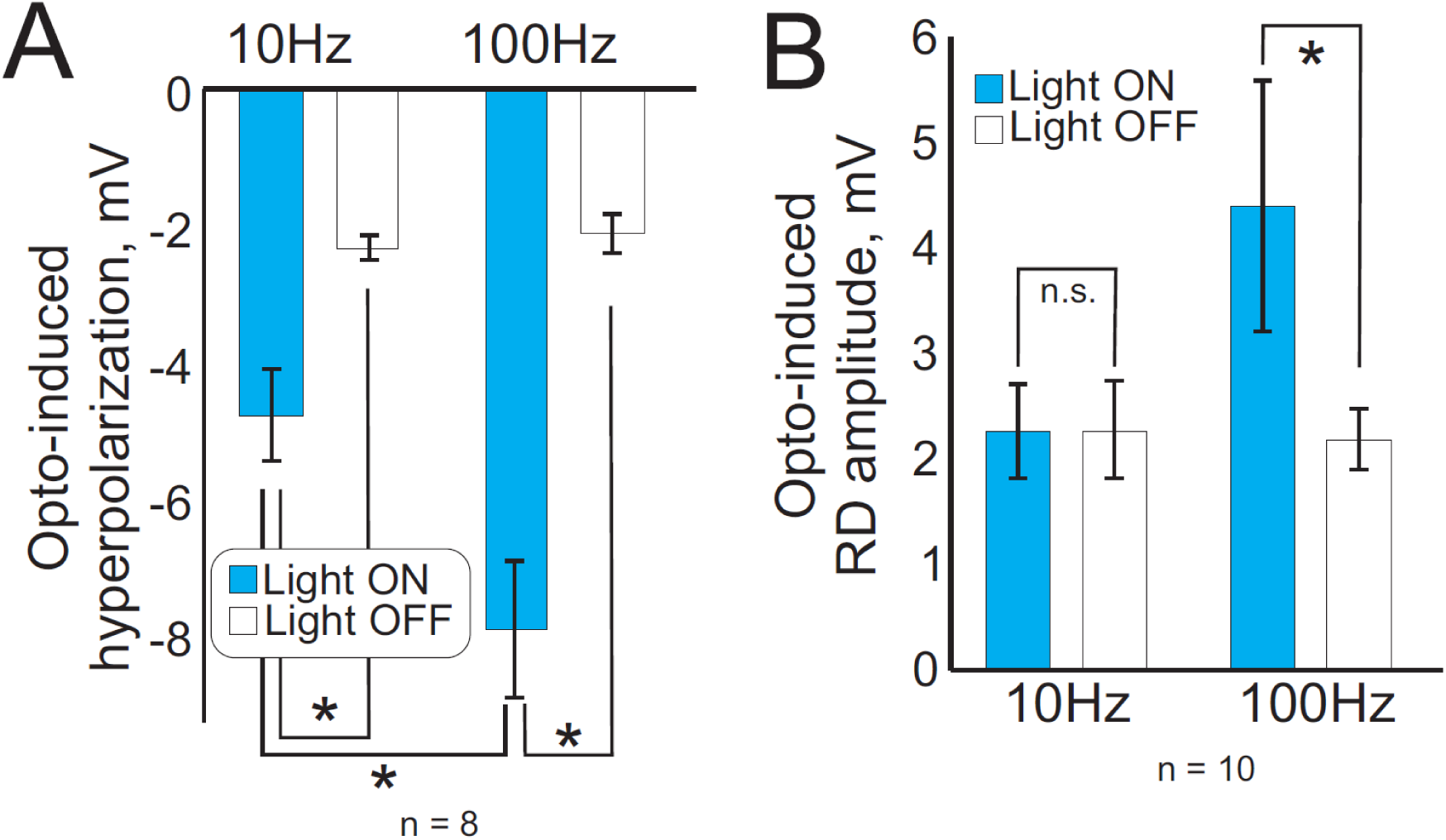
High-frequency stimulation of GABAergic nigral terminals in the SC evokes hyperpolarization and RD in PDB neurons. In Figure 4 of the main text, we show that higher frequency light stimulation is more likely to evoke RD and RD-evoked spiking in PDB neurons compared to lower frequency stimulation. To assess this quantitatively, we measured the hyperpolarization amplitude in PDB neurons induced by 10Hz and 100Hz light stimulation of the GABAergic nigral terminals. (**a**) Shows the amplitude of the peak hyperpolarization in mV for 10Hz and 100Hz during light ON (cyan) and light OFF (white), measured from eight PBD neurons. During light ON, both frequencies evoked hyperpolarizations statistically different from those obtained during light OFF, with 100Hz stimulation on average resulting in larger hyperpolarizations than 10Hz stimulation (n = 8; * p<0.05, paired t-test). (**b**) Shows the same quantification for the amplitude of the RD. During light ON, only 100Hz light stimulation resulted in RD amplitudes statistically different from the RD recorded during light OFF (n = 10; * p<0.05, paired t-test). Cyan bars are with light and white bars are without light stimulation. These results are consistent with the results shown in Figure 4 of the main text wherein we show that 100Hz but not 10Hz light stimulation of Chronos-expressing GABAergic nigral afferents in the SC results in reliable contralateral turning *in vivo*. Referring to Figures 3 and 4 of the main text.

**Supplemental Figure 4.**
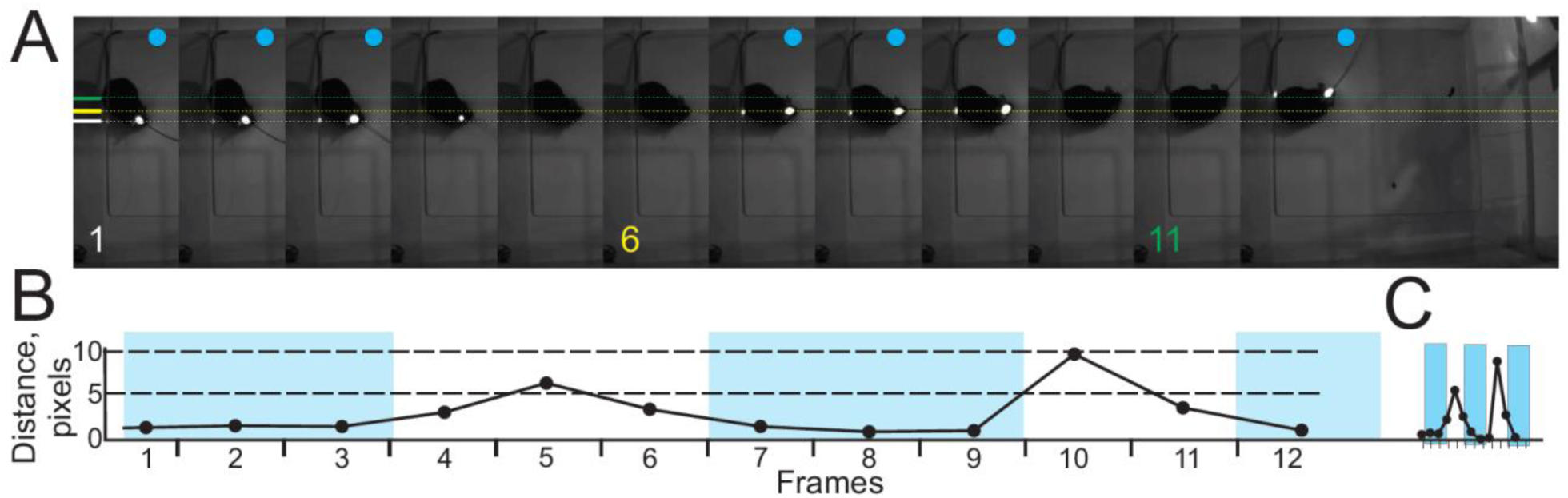
Orienting movements occur at the offset of the optogenetic activation of GABAergic nigra terminals in the SC. Based on our finding that hyperpolarization of PDB neurons from nigral afferents produces RD and RD-evoked spikes *in vitro* (Figure 4), and the unexpected finding that high frequency optogenetic activation of nigral afferents in the SC produces contralateral orienting behavior *in vivo* (Figure 1), we hypothesized that it was the post inhibitory rebound activation of the PDB neurons that produced the orienting behavior. To test this hypothesis, we examined videos on a frame-by-frame basis with light pulse onsets and offsets aligned, to determine when the movement started relative to the onset and offset of the light stimulation. (**a**) Sequence of ∼4 sec video recording during a behavioral session in the open field. Each image shows a single video fame with the frame number indicated at the bottom (1, 6 and 11). Each frame is ∼30ms apart. The cyan circle in each frame shows when the light stimulation was ON. The dotted lines mark the head position at the frames indicated by the color (white 1^st^, yellow 6^th^ and green 11^th^). Optogenetic stimulation occurred in the right SC. The mouse begins to turn its head contralaterally to the site of stimulation within one frame from the offset of the light stimulation. (**b**) Head position distance in pixels (black circles) measured for each consecutive frame during light stimulation (cyan shading) and no stimulation (white). (**c**) A time-compressed representation of the plot shown in (b). Referring to Figures 3 and 4 of the main text.

**Supplemental Figure 5.**
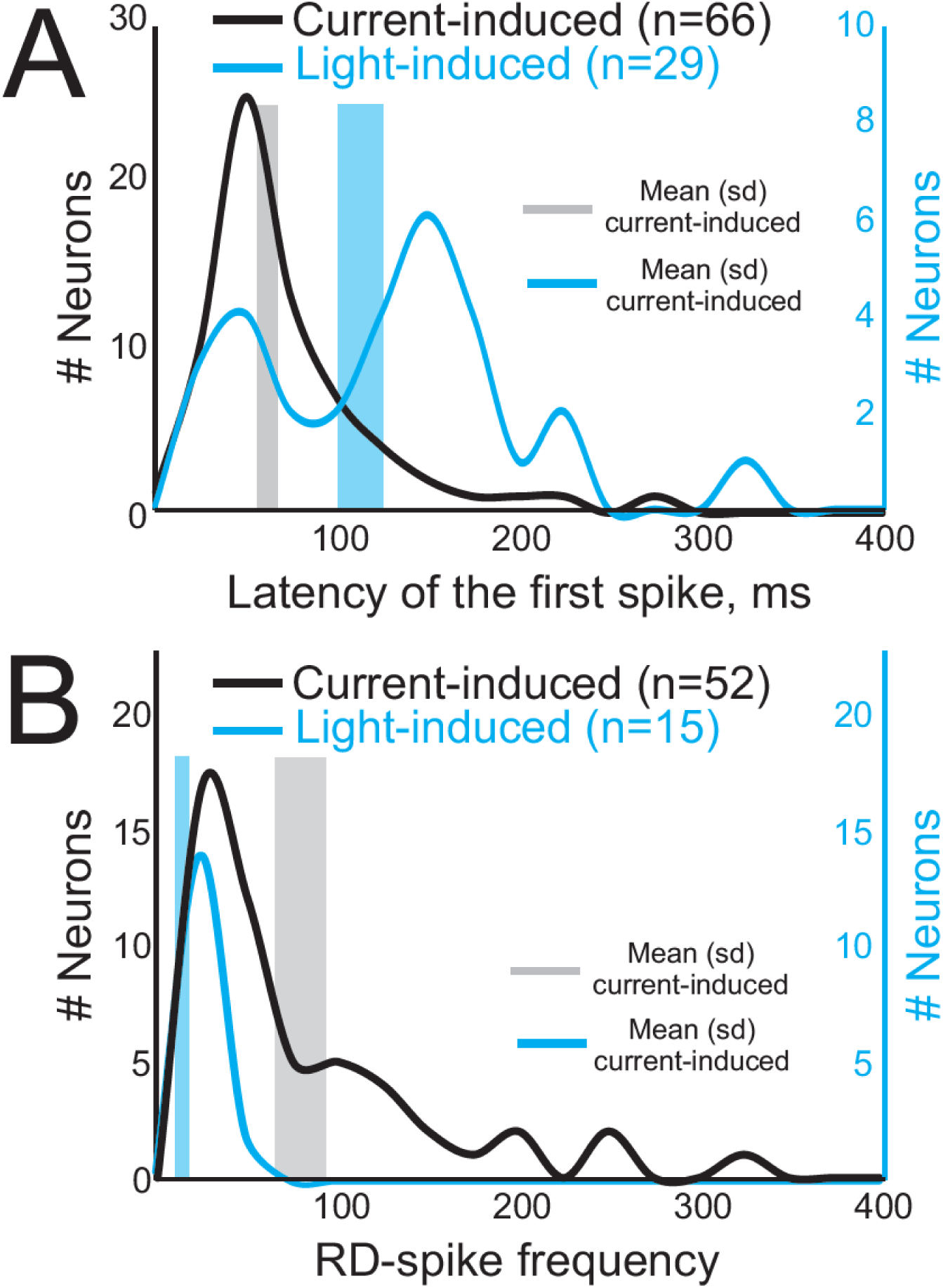
The time-course of RD-evoked spikes in PDB neurons is consistent with the time-course of behavior. In Figures 3 and 4 of the main text, we showed that optogenetic activation of the nigral afferents in the SC produced RD and RD-evoked spiking in PDB neurons of the SC. The same frequency of stimulation also produced orienting behavior *in vivo*. Moreover, the results shown in Supplemental Figure 4 revealed that the orienting movement evoked *in vivo* occurred at the end of the light pulse, consistent with a causal role for the RD and RD-evoked spikes in driving this behavior. Based on this we reasoned that if the RD-evoked spikes in PDB neurons were driving the orienting behavior, then the time-course of these spikes should be similar to those seen in behavior. To assess this, we measured the latency to the first RD-evoked spike and the frequency of the RD-evoked spiking with respect to the end of the hyperpolarization. (**a**) Black lines and left Y-axis show the first spike latency values from neurons where the RD-evoked spikes were triggered by a 500ms duration, −100pA current injection into the PDB neurons. The median latency from the end of the hyperpolarization was 45.85ms and the mean was 77.55ms (SD 13.03; grey vertical bar; bar width shows 1 SE). Most of the neurons showed a latency of 50ms as indicated by the peak of the black curve. The blue lines and the right Y-axis show the latency from the end of the hyperpolarization to the first RD-evoked spikes triggered by 500ms duration, 100Hz light stimulation. The median latency was 120.65ms. The mean latency was 112.78ms (SD 12.86; blue vertical bar; bar width shows 1 SE). The first peak on the blue curve is at 50ms showing that a number of PDB neurons discharge RD-evoked spikes in time sufficient to influence behavior. (**b**) The black line and left Y-axis show the distribution of spike frequencies measured from PDB neurons triggered by 500ms duration, −100pA current injection. The median spike frequency was 49.90 sp/sec and the mean and 1SD indicated by the grey bar and the width of the grey bar respectively, were 77.60 sp/sec and 13.03 sp/sec. The peak of the distribution shown by the black line is at 32.80 sp/sec. The cyan line and right Y-axis show the same for RD-evoked spikes triggered by 500ms duration, 100Hz light pulses. The median spike frequency was 12.20 sp/sec. The mean and 1 SD spike frequency indicated by the blue bar and its width were 13.50 sp/sec and 1.9 sp/sec. The peak of the distribution occurred at 30.5Hz. These measured spike rates are consistent with reported spiking rates for SC neurons *in vivo* in rodents during orienting in spatial choice tasks (e.g., 5-60 sp/sec(Felsen and Mainen, 2008). Referring to Figures 4 and 5 of the main text.

**Supplemental Figure 6.**
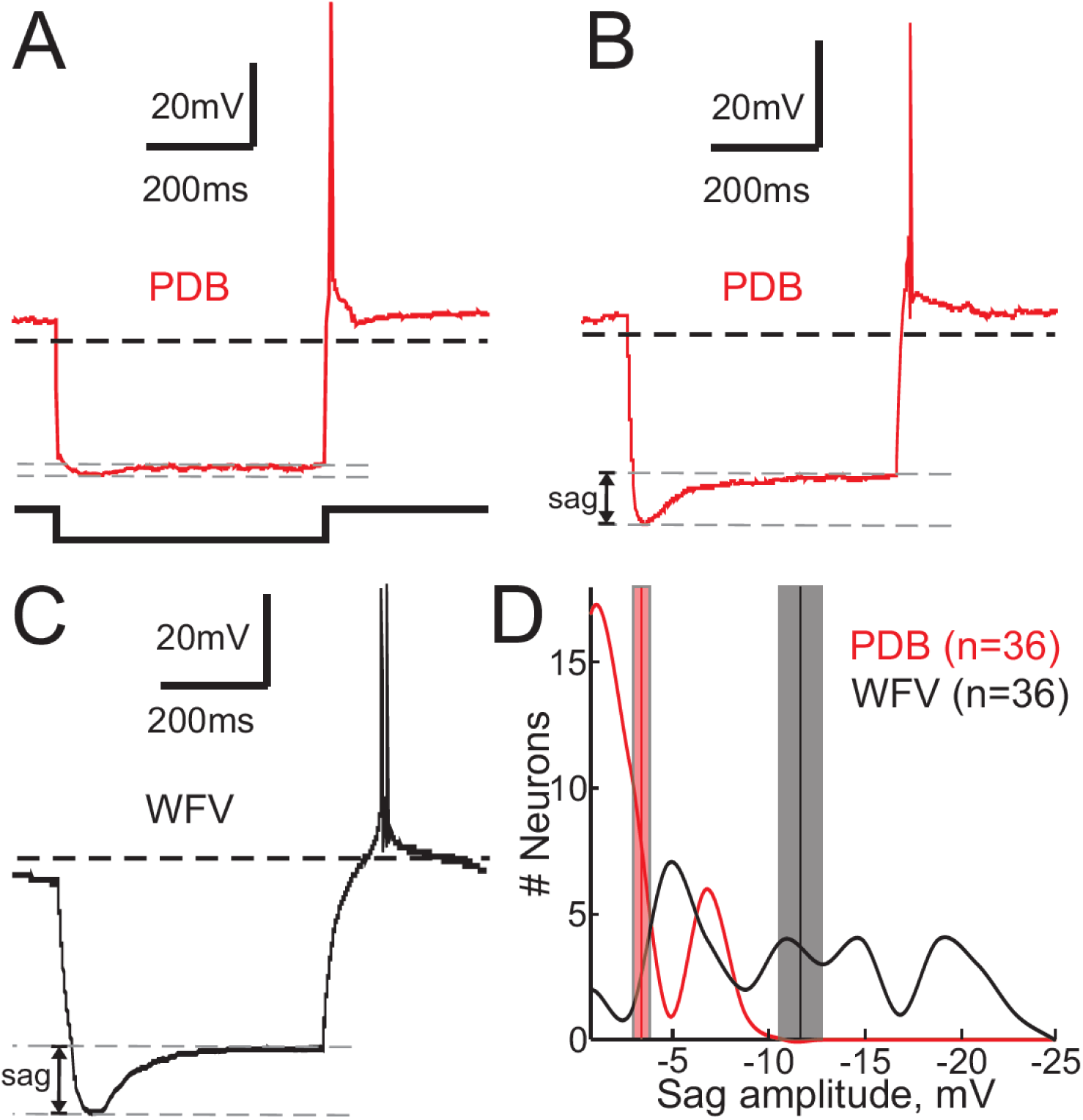
A small proportion of PDB neurons show voltage sag consistent with HCN channel activation. To determine the nature of the channels underlying the RD and RD-spikes in PDB neurons, we introduced negative step currents into PBD neurons and measured the presence or absence of a voltage sag – a defining feature of I_h_. I_h_ is thought to be instrumental in the development of the RD and RD-spikes. We found that most PDB neurons showed very small to no sag voltage (**a**, red trace) upon injection of negative step currents (black trace). Only a small fraction of PDB neurons showed sag voltage with negative step current steps 8/37 (22%). (**b**) On the other hand, most wide-field vertical (WFV) neurons in the superficial layer of the SC present sag voltages with the same step current injections (**c**, black trace). The dashed gray lines show the peak of the sag voltage and the voltage amplitude at the end of the hyperpolarization after I_h_ inactivation. The black dashed lines indicate V_m_ = −60mV. Note that the kinetic of the RD in WFV is slower compared to PDB neurons, concordant with the voltage-dependent kinetic activation of activation of T-type Ca^++^ channels. (**d**) The frequency histogram of the numbers of neurons vs the amplitude of the sag voltage in WFV (black line) and PDB neurons (red line). The vertical lines show the mean sag amplitude value and the shaded rectangle the SE.

**Supplemental Figure 7.**
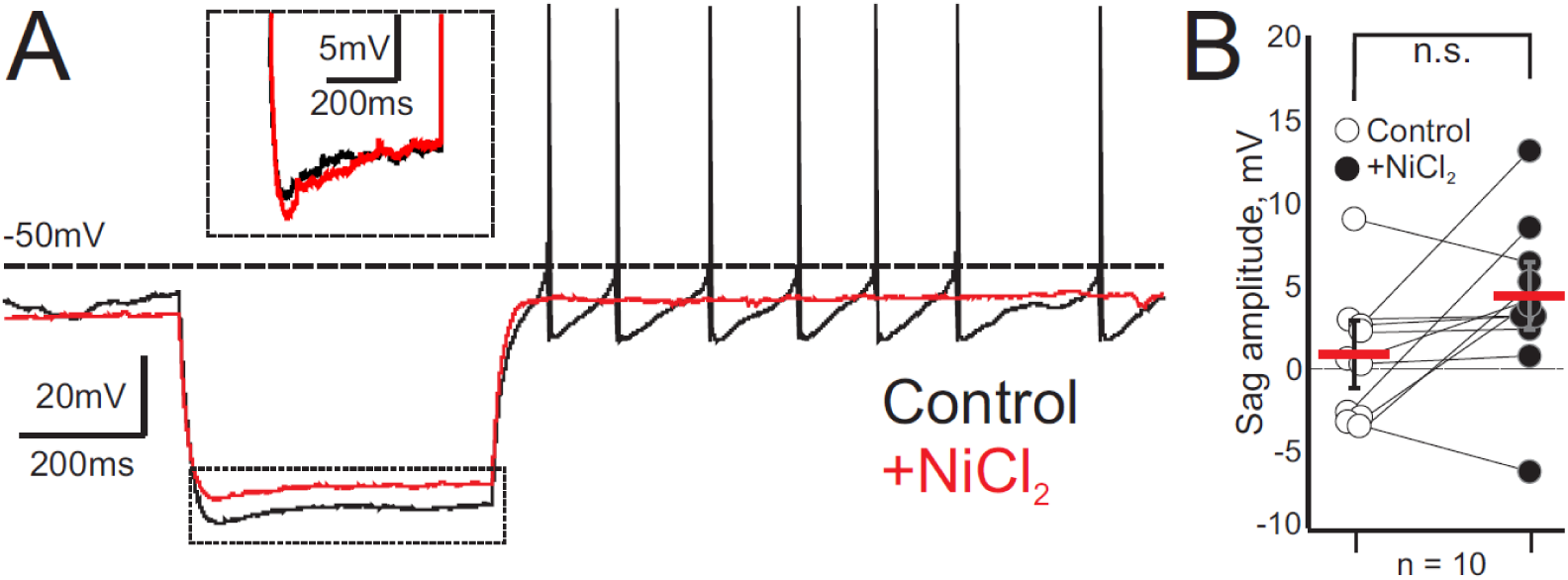
The T-type Ca^++^ channel blocker NiCl_2_ does not inhibit the voltage sag in PDB neurons. Although few PDB neurons showed a voltage sag, when they did it was smaller in amplitude compared to that seen in other neurons including WFV neurons of the superficial SC (Supplemental Figure 6). Nevertheless, in PDB neurons with the sag, we assessed whether the I_h_ was impacted by the T-type Ca^++^ channel blocker NiCl_2_. (**a**) Current clamp traces showing the changes in voltage after a 500ms duration, −100pA current step during control (black trace) and after application of the T-type Ca^++^ channel blocker NiCl_2_ (500μM, red trace). The inset represents a time-compressed and amplitude expanded, portion of the traces outlined by the dashed boxed. Traces were adjusted to compensate for the change in membrane resistance caused by NiCl_2_. (**b**) Sag (n=10) in mV induced by the injection of 500ms duration −100pA current in PDB neurons in control (open circles) and during application of NiCl_2_ (black circles). Red bars, mean ± SE.

**Supplemental Video 1. Optogenetic stimulation of the GABAergic nigro-collicular circuit evokes contralateral orienting movement.** The video shows 10 sec of video at 2X speed obtained from a behavioral session in the open field recorded with a zenithal camera of a GAD2 mouse injected with Chronos in the nigra. Optogenetic stimulation occurred through an optic fiber probe implanted into the lateral iSC on the right side. The video shows ∼2.0 sec of baseline recording where the mouse shows a mixture of random movements and grooming behavior. After the light stimulation (100Hz), the mouse shows a series of continuous turning movements contralateral to the stimulation side. Referring to Figure 1 of the main text.

**Supplemental Video 2.** The video shows 10 sec of video at 2X speed obtained from a behavioral session in the open field of a mouse injected with the control virus (*i.e.*, no Chronos) in the right nigra and with light stimulation of the terminals in the right SC. The video shows the mouse exhibiting no systematic movements before or after the light stimulation of the nigral afferents in the right SC. Referring to Figure 1 of the main text.

**Supplemental Video 3.** The video shows a 37 second video clip at 0.12x speed, from an open field behavioral measurement session showing the orienting movement evoked by light stimulation of the right SC in a GAD2 mouse injected with Chronos in the right nigra. The light stimulation was of the terminal located in the right SC. The slowing of the video shows that the initiation of the orienting movement occurs around the time that the light pulses end rather than with the initiation of the optogenetic stimulation. The video shows a series of 17 pulses. Each light pulse corresponds to a 100ms/100Hz light pulse stimulation. Referring to Figure 4c-d of the main text.

## Notes

### Competing Interest Statement

The authors have declared no competing interest.

